# Eco-evolutionary dynamics and environmental detoxification jointly shape bacterial community response to antibiotic perturbation

**DOI:** 10.64898/2026.04.27.721019

**Authors:** Johannes Cairns, Niina Smolander, Sanna Pausio, Olli Pitkänen, Meri Lindqvist, Manu Tamminen, Rishi Das Roy, Ville-Petri Friman, Lutz Becks, Ville Mustonen, Teppo Hiltunen

## Abstract

Microbial communities frequently encounter recurrent antibiotic disturbance, yet how ecological, evolutionary, and environmental processes jointly shape the response remains unresolved. Here we experimentally disentangle these mechanisms using a 23-species bacterial community exposed to ampicillin pulses with pre-pulse and resistance priming. Pre-pulse priming preconditioned community composition toward resistant taxa, reducing compositional change during the main pulse. Resistance priming was the dominant determinant of community response, buffering compositional change, relaxing selection, and altering gene expression. This buffering arose from both evolution of higher resistance and accelerated ampicillin detoxification by a dominant degrader, which transiently reduced antibiotic effects and favoured non-degrading taxa. Yet buffering did not translate into better recovery: because resistance was coupled to competitive dominance, diversity remained comparable to or below that of ancestral communities, and dominance was reinforced. Together, we show that antibiotic history reshapes microbial disturbance response through coupled eco-evolutionary and environmental feedbacks, generating a trade-off between resistance and recovery.

## Introduction

Understanding how microbial communities respond to disturbances is fundamental for medicine, biotechnology, and ecosystem management [1–3]. Community responses are typically attributed to ecological interactions among species and evolutionary changes within them [4], yet the abiotic environment can shift dynamically during disturbance and feed back on both. Although resource-consumer theory captures environmental feedbacks on ecological and evolutionary dynamics [5, 6], empirical work explicitly integrating all three processes remains limited, particularly for microbial communities comprising more than two species.

Such communities underpin essential ecosystem functions but are sensitive to anthropogenic stressors such as antibiotics, heavy metals, and agrochemicals [4, 7–10]. The community response to disturbance can be decomposed into resistance, the capacity to remain unaltered during disturbance, and recovery (or resilience), the capacity to return toward the pre-disturbance state [4]. Because microbes evolve rapidly on ecological timescales owing to their large population sizes and short generation times, both ecological species sorting and rapid evolutionary change can contribute to these responses.

Prior exposure to stress, ‘priming’ [4, 11], can reshape resistance and recovery through interacting ecological and evolutionary mechanisms. At the ecological level, priming filters susceptible taxa and enriches resistant ones, shifting community composition toward states more tolerant of subsequent disturbance. At the evolutionary level, priming can generate increased resistance within taxa, further buffering communities against re-exposure [11, 12]. Recovery will depend on the alignment between stress resistance and competitive ability among taxa that persist through disturbance: if resistant taxa are strong competitors, diversity may remain durably reduced; if resistance carries fitness costs, previously suppressed taxa may re-establish. Despite these predictions, experiments that disentangle ecological, evolutionary, and environmental contributions to priming effects across both disturbance and recovery phases remain scarce [11, 12].

One mechanism capable of coupling environmental change to community dynamics during antibiotic disturbance is enzymatic antibiotic degradation. β-lactamases, which hydrolyse β-lactam antibiotics such as ampicillin, can act as public goods by reducing antibiotic concentrations in the shared environment [10, 13, 14]. Declining concentrations may extend the conditions under which less resistant taxa persist [15], restructure competitive hierarchies [15], and reduce the benefit of resistance [16], thereby shaping subsequent recovery dynamics. Moreover, resistance mechanisms and degradation capacity can evolve rapidly under antibiotic selection [17, 18], and resistance evolution in non-degraders may erode exposure protection [19]. Species interactions can further modulate these evolutionary trajectories [20]. Antibiotic degradation therefore provides a tractable context in which ecological sorting, rapid evolution, and environmental transformation interact through coupled feedbacks.

Our previous work using a 23-species synthetic bacterial community showed that short antibiotic pulses produce highly repeatable community trajectories largely explained by intrinsic species traits, namely growth rate and antibiotic susceptibility [21]. We have also shown that disturbance history shapes subsequent responses through both ecological and evolutionary dynamics [12, 22]. However, whether community-driven modification of the antibiotic environment feeds back on these eco-evolutionary processes has not been explicitly tested.

Here, we assembled four otherwise identical 23-species communities differing only in species-level resistance priming history and subjected them to a two-pulse ampicillin perturbation experiment varying pre-pulse intensity (Fig. 1). We predicted that species-level resistance priming would increase community resistance and reduce the importance of community-level pre-pulse priming [11, 12], whereas ancestral communities would depend more strongly on ecological sorting and rapid resistance evolution during pre-pulse exposure [3, 21, 23, 24]. We further predicted that priming could alter antibiotic degradation dynamics through ecological or evolutionary changes, thereby modifying antibiotic exposure and amplifying community-level protection [15]. By quantifying community composition, functional traits, antibiotic degradation dynamics, and genomic and transcriptional responses, we disentangle how ecological filtering, evolutionary priming, and community-driven environmental change shape microbial resistance and recovery under recurrent antibiotic disturbance.

**Figure 1.**
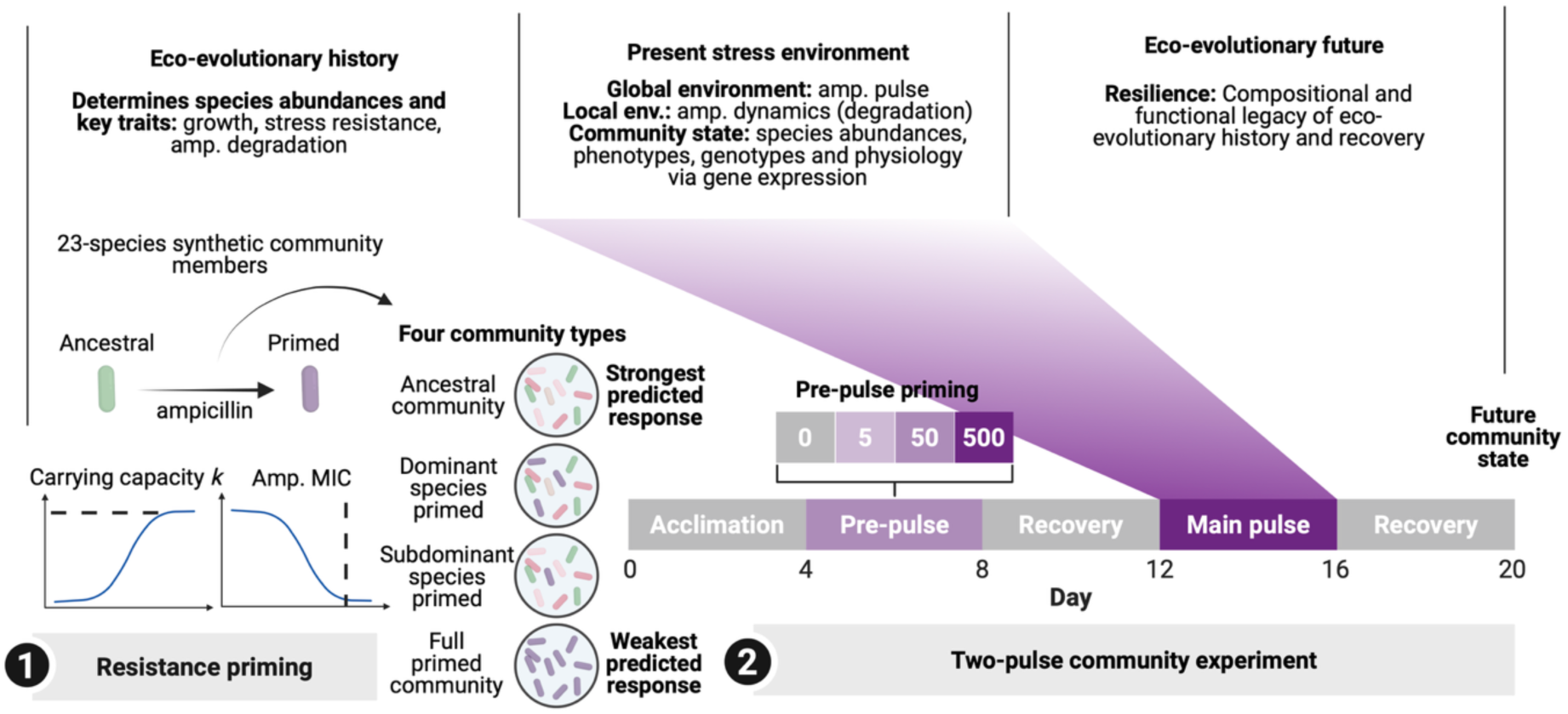
Experimental design and framework. Schematic overview of the experimental design linking species-level resistance priming histories, community-level antibiotic pulses, and disturbance response trajectories. **(1) Resistance priming:** Each of the 23 bacterial species was first cultured at increasing ampicillin concentrations to generate paired ancestral and primed strains differing in carrying capacity (*k*) and ampicillin resistance (MIC). These strain sets were assembled into four community types differing only in the priming history of their members: ancestral community; dominant species *Aeromonas caviae* primed community; subdominant species *Pseudomonas chlororaphis* primed community; full primed community. **(2) Two-pulse community experiment:** Communities were propagated through a 20-day serial-transfer regime comprising five 96 h epochs: acclimation, pre-pulse priming, intermediate recovery, main pulse, and final recovery. Pre-pulse ampicillin concentrations were 0, 5, 50, or 500 µg mL^−1^; all communities received 500 µg mL^−1^ during the main pulse. We quantified responses across species traits, community composition, genomic and transcriptional changes, and antibiotic degradation dynamics. **Alt text:** Two-panel schematic. Panel 1 shows resistance priming: 23 species are exposed to escalating ampicillin to generate ancestral and primed strain pairs, assembled into four community types. Panel 2 shows the two-pulse experiment timeline with five labelled phases across 20 days, with pre-pulse and main pulse concentrations indicated.

## Materials and Methods

### Model community and culture conditions

The study employed a previously established 23-species gram-negative synthetic bacterial community [12, 21, 25, 26]. Community members, isolated from diverse habitats (soil, water, or host organisms), were selected for their ability to grow in uniform culture conditions and for bioinformatic separability via 16S rRNA marker gene sequences. All species have been phenotypically characterised and whole-genome sequenced [26, 27]. All experiments used Reasoner’s 2A (R2A) liquid medium at 30 °C with shaking at 1000 rpm, unless stated otherwise.

### Species-level resistance priming

All 23 community members were individually resistance primed through stepwise-escalating ampicillin exposure. Monocultures were grown in 100 % R2A at each species’ IC50 ampicillin concentration at 25 °C with constant shaking (70 rpm), serially transferred every 96 h (8% transfer volume), and exposed to doubling ampicillin concentrations until growth ceased. Single clones were isolated from the highest ampicillin concentration supporting growth and freeze-stored with glycerol at −80 °C.

### Estimating species growth and ampicillin susceptibility

Growth characteristics and ampicillin susceptibility were determined from growth curve measurements (BioTek LogPhase 600 Microbiology Reader) across 23 ampicillin concentrations (0.27–2000 µg mL^−1^, 2:3 dilution series) in two technical replicates per species (100 % R2A, 150 µL, 30 °C, 800 rpm, optical density at 600 nm every 10 min for 48 h). Carrying capacity (*k*) was extracted from fitted growth curves; *k* values below 0.1 or above 1.3 were excluded. The minimum inhibitory concentration (MIC) was the lowest ampicillin concentration at which no growth was observed, and MIC values were log10-transformed for linear modelling.

### Two-pulse community experiment

Four 23-species communities were assembled differing only in resistance priming history: (i) ancestral community (all ancestral clones); (ii) dominant species *Aeromonas caviae* primed; (iii) subdominant species *Pseudomonas chlororaphis* primed; and (iv) full primed community (all species primed). *A. caviae* was chosen as the consistent dominant community member under antibiotic-free conditions [12, 21]; *P. chlororaphis* was chosen as an abundant member exhibiting the strongest resistance-associated carrying capacity cost after priming (Fig. 2A).

**Figure 2.**
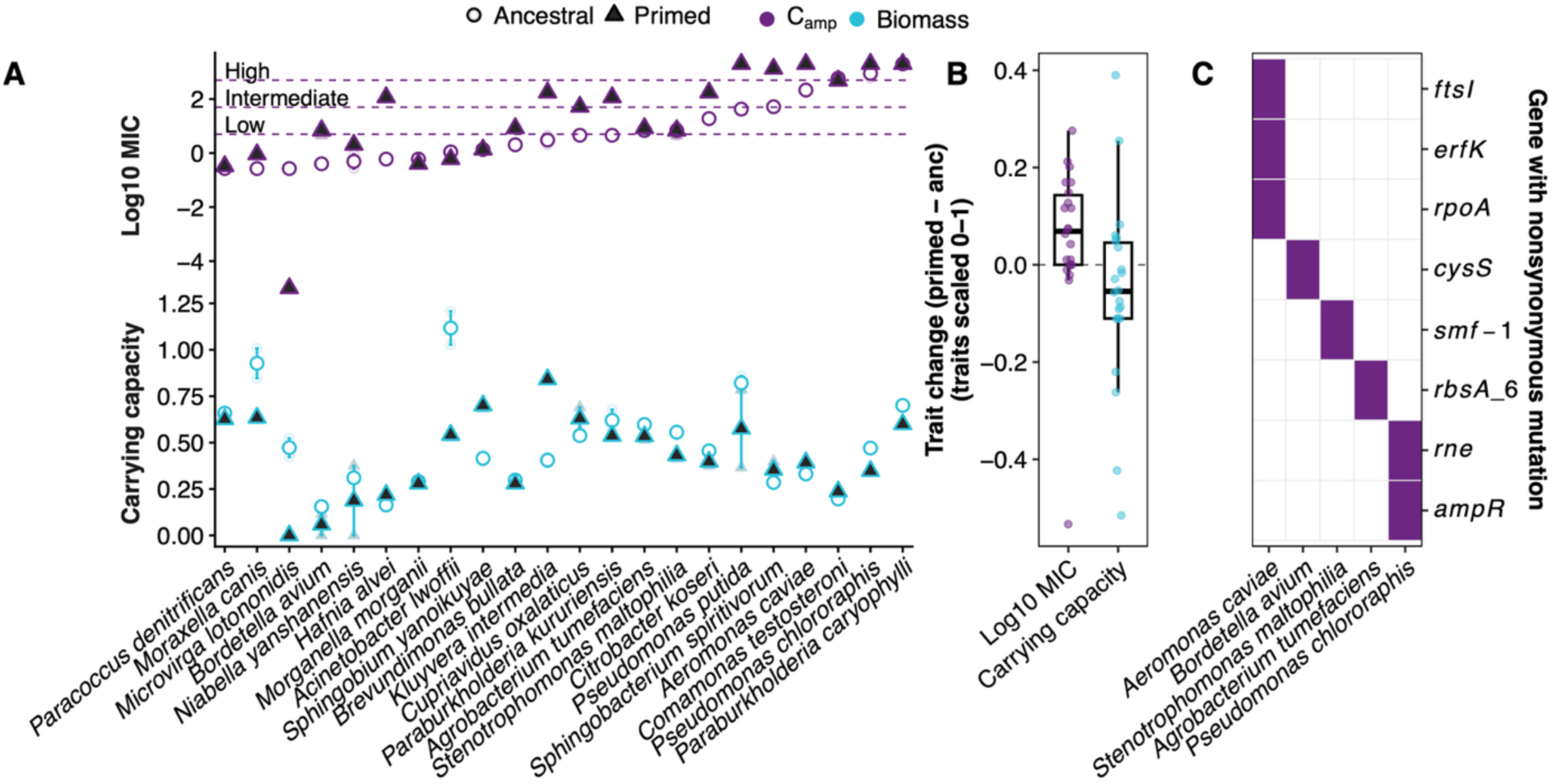
Trait and genomic evolution of resistance primed species. **(A)** Ampicillin resistance (minimum inhibitory concentration, MIC; purple) and carrying capacity (*k*; turquoise) of synthetic community members prior to (circles) and after (triangles) individual resistance priming. Data show mean and bootstrapped 95 % confidence intervals (n = 2 technical replicates per clone). **(B)** Change in MIC and carrying capacity (traits scaled 0–1) between resistance primed and ancestral strains (mean ± s.e.m.) across species. **(C)** Genes with nonsynonymous mutations in resistance primed clones representing abundant community members. **Alt text:** Three-panel figure. Panel A shows a dot-and-triangle plot for each of 23 species, with purple symbols for MIC and turquoise for carrying capacity before and after priming; most species show increased MIC and decreased carrying capacity after priming. Panel B shows a box plot of mean change in MIC versus change in carrying capacity across species, with most species increasing in MIC and decreasing in carrying capacity. Panel C shows a heatmap of nonsynonymous mutations across species and genes, with cells coloured by mutation presence.

A full-factorial serial passage experiment crossed four resistance priming histories with four community-level pre-pulse ampicillin concentrations (0, 5, 50, or 500 µg mL^−1^) in four biological replicates each, yielding 64 communities in total. Communities were passaged through five 96 h epochs: acclimation, ampicillin pre-pulse, intermediate recovery, main pulse (500 µg mL^−1^ for all communities regardless of pre-pulse level), and final recovery, with serial transfer every 24 h (10 % transfer volume) into 48-well plates (2.5 mL total volume). Cell density was estimated by flow cytometry (Guava easyCyte 5 HT HP, Cytek) every 24 h. Samples for 16S rRNA amplicon sequencing, liquid chromatography–mass spectrometry (LC-MS) analysis of ampicillin concentration, and freeze-storage were collected every 96 h (end of each epoch). RNA-seq and metagenomic samples were collected after the main pulse (day 16) and at experiment end (day 20), respectively. Communities were assembled from monoculture stocks equalised to 5 × 10^6^ cells mL^−1^ in M9 salt solution by flow cytometry-estimated density, combining 250 µL per species.

### Mass spectrometric analysis of ampicillin degradation

Ampicillin degradation was assessed by monoculture LC-MS assays on four species that grew or persisted during the main community pulse (*Aeromonas caviae*, *Bordetella avium*, *Comamonas testosteroni*, and *Pseudomonas chlororaphis*) and by leave-one-out community experiments that sequentially removed each of the three identified degraders, or all three simultaneously, from ancestral or full primed community backgrounds. Cultures were inoculated at 500 µg mL^−1^ ampicillin in 1 mL R2A; cell-free supernatants were collected at 6, 12, and 24 h by centrifugation and 0.2 µm filtration and stored at −20 °C. Ampicillin concentrations were quantified by triple-quadrupole LC-MS with deuterated ampicillin as internal standard (Agilent 6460, Turku Metabolomics Centre).

### DNA extraction and sequencing

Genomic DNA was extracted using the DNeasy 96 Blood & Tissue Kit (Qiagen) following the manufacturer’s protocol. 16S rRNA amplicon libraries (V3 region; TaggiMatrix system [28, 29]) were sequenced on an Illumina MiSeq platform (2 × 150 bp). Whole-genome sequencing (WGS) of resistance primed clones from 10 dominant species was performed using Illumina DNA Prep kits (SeqCenter, https://www.seqcenter.com; 2 × 151 bp, NovaSeq X Plus). Metagenome and RNA-seq libraries were sequenced on an Illumina NovaSeq 6000 (Finnish Functional Genomics Centre, FFGC). RNA was extracted using a Monarch Total RNA Miniprep Kit (New England Biolabs), with bead-beating disruption in a TissueLyser II (Qiagen) and on-column DNase treatment.

### Sequence data processing

16S rRNA amplicon data were processed by demultiplexing, quality trimming, paired-end merging, and mapping to community reference sequences [30], with normalisation by 16S gene copy number per species. WGS and metagenomic variants were identified by aligning reads to a custom HAMBI reference genome database (BWA-MEM [31]), followed by GATK Mutect2 [32] variant calling, hard filtering of single nucleotide polymorphisms (SNPs) and insertions/deletions (INDELs), SnpEff [33] annotation, and retention of protein-altering changes only [12]. Copy number variants (CNVs) were detected using CNproScan v0.999 with GC content and mappability normalization [34]; recurrent duplications (≥2 independent communities) were retained. RNA-seq reads were pseudoaligned using Salmon v1.10.1 [35] via the nf-core rnaseq pipeline v3.14.0 [36], with community-composition-based normalisation of expression values [25]; differential expression was identified using DESeq2 v1.44.0 [37] (false discovery rate [FDR] < 0.05, |log2 fold change| > 1). Full bioinformatics parameters, quality filters, and variant calling thresholds are provided in the Supplementary Methods.

### Statistical analysis

All analyses were conducted in R v4.4.2 [38]. Community composition was analysed using Bray-Curtis dissimilarity and permutational multivariate analysis of variance (PERMANOVA; adonis2 in vegan [39]; 999 permutations, type = ‘by terms’). Temporal instability was quantified as trajectory roughness (cumulative |ΔBray-Curtis| across days 4–20).

Community-level traits were computed as scalar products between species phenotypes and relative abundance in the community (*f*) with generalised least squares (GLS) models (nlme [92]; first-order autoregressive autocorrelation structure). Trait values for the ancestral or resistance primed species were used depending on which species was present in the corresponding community treatment. For a treatment panel, at day *t*:

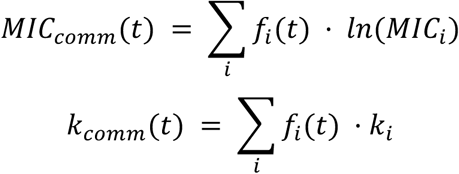

The effects of experimental treatments on cell density, species richness, Shannon diversity, community MIC, community carrying capacity (*k*), and ampicillin degradation dynamics were analyzed using generalized least squares (GLS) models (nlme [40]; first-order autoregressive autocorrelation structure). Species MIC and *k* were compared using ANOVA with species × priming interaction terms. Ampicillin degradation kinetics were analysed using log-rank tests and restricted mean survival time (RMST) to 24 h. Pairwise contrasts were performed with emmeans [41], and model selection with stepAIC in MASS [42]. MIC and carrying capacity (*k*) of ancestral vs. resistance primed species were assessed with linear regression, and mutation counts with Poisson regression. For species compositional ordination plots, t-SNE was applied to the species-by-sample abundance matrix using Rtsne [43, 44] (perplexity = 15, θ = 0.5, dims = 2, PCA preprocessing enabled). Portions of the analysis code were refactored with the assistance of ChatGPT (GPT-5.2 Thinking; OpenAI, accessed February 2026). All code and outputs were verified by the authors.

## Results

### Resistance priming drove resistance-growth trade-offs

Resistance priming broadly increased MIC levels across the 23 community members, with marked species-specific variation in magnitude (ANOVA on log10 MIC: resistance priming *F*1,46 = 28.41, *p* < 0.001; species × priming *F*22,46 = 6.73, *p* < 0.001; Supplementary Table S1; Fig. 2A,B): 19 of 23 species exhibited higher MIC after priming. Evolving resistance imposed a fitness cost in carrying capacity (*k*), with primed species showing decreased *k* in antibiotic-free conditions, contingent on species identity (ANOVA: priming *F*1,46 = 4.97, *p* = 0.031; species × priming *F*22,46 = 5.29, *p* < 0.001; Supplementary Table S2; Fig. 2A,B): 15 of 23 species showed reduced *k*. Despite differences between species, MIC change also correlated positively with change in *k* across species (Spearman’s ρ = 0.47, *p* = 0.025). These results are consistent with classical resistance-fitness trade-offs [45, 46].

Whole-genome sequencing of 10 dominant resistance primed species identified fixed nonsynonymous mutations or large structural variants in six (Fig. 2C; Supplementary Results). In *A. caviae*, mutations occurred in *ftsI* (encoding penicillin-binding protein 3 [PBP3]; involved in peptidoglycan cross-linking and β-lactam resistance [28,29]) and *erfK* [29], together with a mutation in *rpoA* (RNA polymerase complex; linked to β-lactamase-independent high-level resistance [30]). *Sphingobacterium spiritivorum* showed a 13× amplification of a genomic block containing the β-lactamase gene blaB [31]. *P. chlororaphis*, which exhibited the strongest *k* decrease after priming, carried mutations in *rne* [35] (potentially reducing OmpA porin abundance and conferring resistance via reduced permeability [36]) and *ampR* [37] (a regulator of AmpC β-lactamase production). These and additional species-specific genomic findings are reported in the Supplementary Results. Together, phenotypic and genomic changes generated trait differences expected to shape community dynamics under antibiotic disturbance.

### Priming type determined resistance-recovery dynamics by altering trait distribution

The four communities differing in resistance priming history were subjected to the two-pulse experiment and sampled before and after pulses and recovery periods. Increasing pre-pulse ampicillin shifted community composition toward the post-disturbance state prior to the main pulse, indicating ecological pre-conditioning through species sorting (Figs 3, 4A). The magnitude of compositional change during the main pulse depended on resistance priming background: ancestral and single-species-primed communities showed pronounced restructuring, whereas full-primed communities exhibited buffered trajectories consistent with community-wide resistance evolution (interaction: *F*3,56 = 4.36, *p* = 0.008; Supplementary Table S3; Figs 3, 4A).

**Figure 3.**
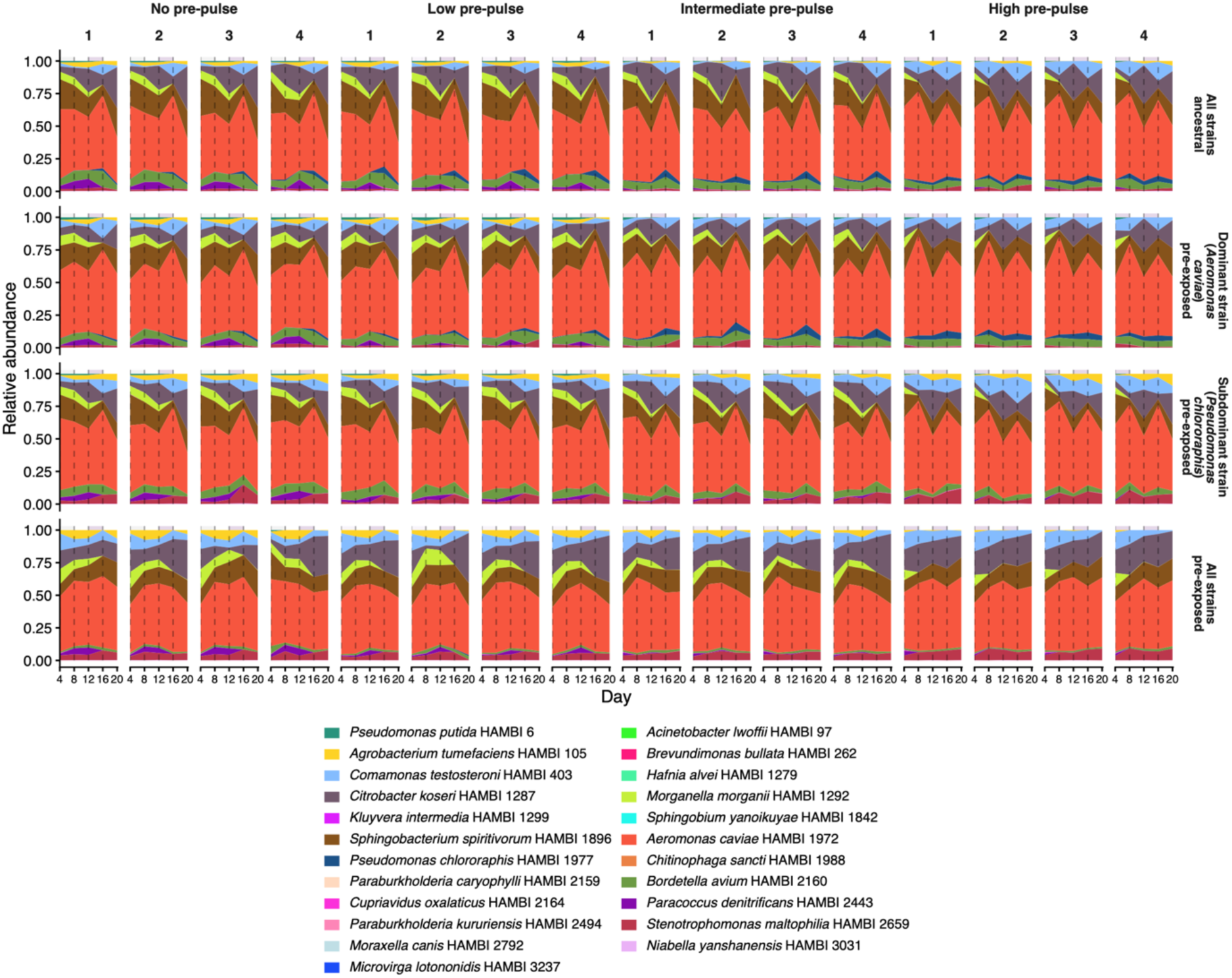
Community composition over the 20-day ampicillin pulse experiment across pre-pulse priming levels and resistance priming backgrounds. Stacked area charts show the relative abundance of constituent community members (colours) across time (sampled on days 4, 8, 12, 16, and 20). Columns indicate pre-pulse ampicillin concentration (µg mL^−1^): 0, 5, 50, and 500. Rows indicate resistance priming background: ancestral community; dominant species (*A. caviae*) primed; subdominant species (*P. chlororaphis*) primed; and full-primed community. Each treatment combination was propagated in four independent biological replicates (n = 4; shown as separate columns within each treatment). Vertical dashed lines mark phase boundaries at days 4, 8, 12, and 16 (pre-pulse phase: days 4–8; main pulse phase: days 12–16). **Alt text:** Grid of stacked area charts with 16 columns (four pre-pulse levels times four replicates) and four rows (resistance priming backgrounds). Each chart shows relative abundance of species across time. Full-primed communities (bottom row) show more stable compositions during the main pulse compared with ancestral communities (top row), which show pronounced compositional shifts, particularly at higher pre-pulse concentrations.

**Figure 4.**
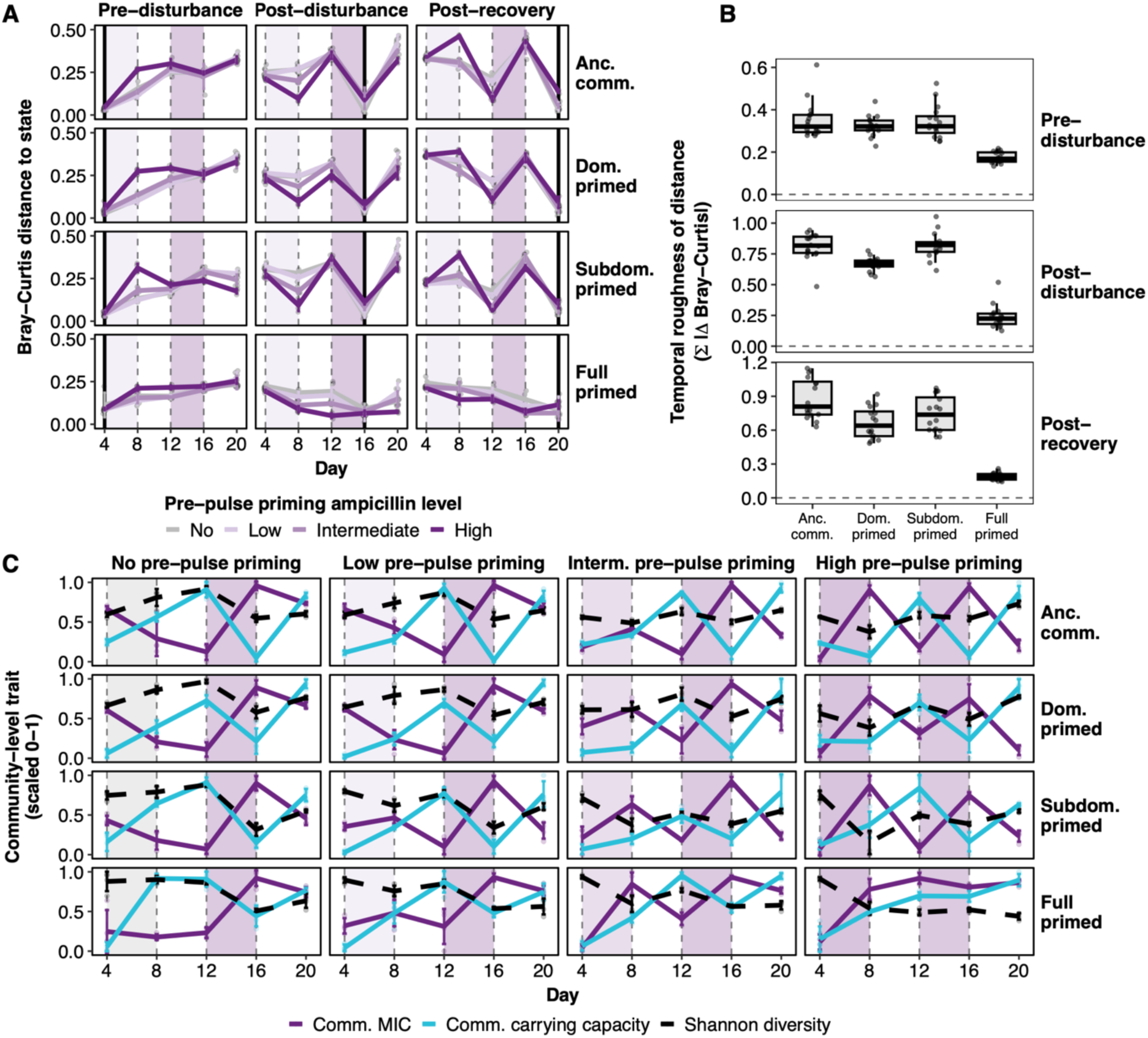
Community dynamics across pre-pulse and resistance priming backgrounds. **(A)** Bray-Curtis dissimilarity of each community from its resistance-priming-background-specific reference centroid at pre-disturbance (day 4), post-disturbance (day 16), and post-recovery (day 20), indicated by black vertical lines. Points are individual replicates (n = 4 per treatment); solid lines show means with bootstrapped 95 % confidence intervals. Shaded regions denote pre-pulse (days 4–8) and main pulse (days 12–16) phases. Rows correspond to resistance priming backgrounds. **(B)** Temporal roughness (Σ|ΔBray-Curtis| across days 4–20). Box plots show median, interquartile range, and 1.5× interquartile range whiskers (n = 16 per resistance priming background). **(C)** Community-weighted mean MIC (log-scale) and carrying capacity (*k*), and Shannon diversity, across time. Traits were min–max scaled within each pre-pulse × background combination. Points are replicates; solid (MIC, *k*) and dashed (Shannon) lines show means with bootstrapped 95 % confidence intervals. Columns: pre-pulse ampicillin concentrations (0, 5, 50, 500 µg mL^−1^). **Alt text:** Three-panel figure. Panel A shows time-series line plots of Bray-Curtis dissimilarity from reference states for each of four community types across 20 days; full-primed communities show the lowest dissimilarity during the main pulse. Panel B shows box plots of temporal roughness by community type; full-primed communities have markedly lower roughness than other types. Panel C shows time-series plots of community-weighted MIC, carrying capacity, and Shannon diversity across pre-pulse levels and community types; Shannon diversity recovers less in full-primed communities.

These patterns were driven by abundant taxa responding in line with their intrinsic traits. In ancestral communities, *Aeromonas caviae* and *Comamonas testosteroni* increased during pulses while *Citrobacter koseri*, *Morganella morganii*, and *Sphingobacterium spiritivorum* declined, consistent with their respective MIC levels (Fig. 2A). In the full primed community, evolved resistance in *C. koseri* and *S. spiritivorum* reduced their decline or promoted their increase, while resistance-primed *P. chlororaphis*, despite high MIC, went near-extinct owing to its strong carrying capacity cost (Supplementary Fig. S1). Cell densities were comparable across treatments (Supplementary Fig. S2), confirming relative abundance changes broadly reflected absolute dynamics.

Quantitative community-level analyses confirmed these patterns. Temporal instability was lowest in full-primed communities across all phases (ANOVA, all *p* < 0.001; Fig. 4B; Supplementary Table S4), with post-disturbance roughness markedly reduced (0.19 ± 0.03 vs. 0.67–0.88 in other treatments), demonstrating enhanced ecological resistance to perturbation. Higher pre-pulse intensity shifted communities toward higher community MIC and lower *k* prior to the main pulse (*F*1,280 = 103.0, *p* < 0.001 and *F*1,280 = 12.2, *p* < 0.001, respectively; Supplementary Tables S5–S6), and stronger pre-pulses advanced community proximity to the final post-recovery state already during intermediate recovery (*F*1,56 = 51.6, *p* < 0.001; Supplementary Table S7), indicating that pre-pulse priming effectively advanced recovery dynamics in time. These trait shifts were largely independent of resistance priming background.

Critically, resistance priming came at a recovery cost. In full-primed communities, increasing pre-pulse intensity reduced species richness (*F*1,307 = 40.19, *p* < 0.001; Supplementary Fig. S3; Supplementary Table S8) and limited recovery of Shannon diversity (*F*3,295 = 26.68, *p* < 0.001; Fig. 4C; Supplementary Table S9). These effects arose from dominance of taxa such as *C. koseri* and *S. spiritivorum*, which had evolved moderate to high resistance with low fitness cost. Thus, resistance priming buffered communities during disturbance but constrained recovery by reinforcing competitive dominance among resistant taxa.

### Environmental feedbacks: resistance priming of dominant degrader accelerated ampicillin breakdown and protected non-degraders

Ampicillin concentrations declined to near-zero within each 24 h transfer cycle across all treatments, with slightly faster degradation at higher pre-pulse levels (pre-pulse × day: *F*1,304 = 17.52, *p* < 0.001; Supplementary Fig. S4; Supplementary Table S10), consistent with ecological enrichment of degraders. Because community-level degradation was nearly complete within each cycle, we focused on within-cycle dynamics to identify key degraders and assess the effects of resistance priming on detoxification rate.

Monoculture LC-MS assays identified *Aeromonas caviae*, *Comamonas testosteroni*, and *Pseudomonas chlororaphis* as rapid ampicillin degraders (Fig. 5A). Resistance priming specifically accelerated ampicillin degradation in *A. caviae*: primed clones reached below 10 µg mL^−1^ by 6 h rather than 12 h (Wilcoxon *W* = 0, *p* = 0.013), with higher biomass at 6 h (OD600: 0.23 vs. 0.04; *p* < 0.001). Degradation timing correlated negatively with growth rate at 6 h (Spearman’s *ρ* = −0.88, *p* = 0.021), indicating that accelerated detoxification arose from improved growth under antibiotic stress rather than intrinsically enhanced enzymatic capacity. By contrast, resistance priming did not shift degradation timing in *C. testosteroni*, while primed *P. chlororaphis* showed reduced early biomass consistent with its growth cost rather than any improvement in detoxification.

**Figure 5.**
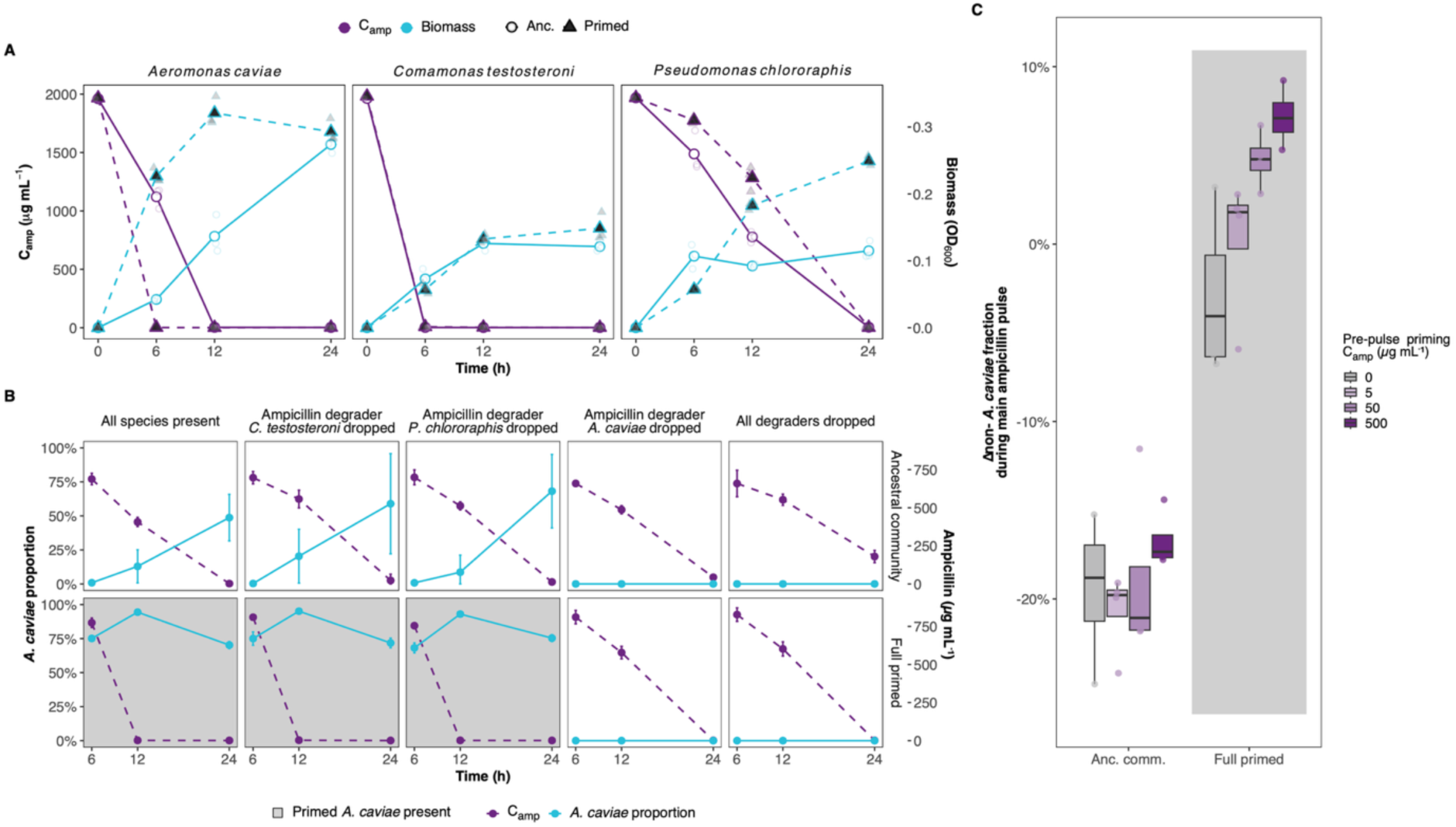
***Aeromonas caviae* dynamics and community responses to ampicillin in monoculture, leave-one-out, and main community experiments.** (A) Ampicillin degradation by ancestral (purple) and resistance primed (turquoise) clones of species that persisted during the main pulse. Data show mean and bootstrapped 95 % confidence intervals (n = 3 technical replicates). (B) *A. caviae* relative abundance (left y-axis, red line) and ampicillin concentration (right y-axis, purple dashed line) at 6, 12, and 24 h in a 24 h leave-one-out experiment (n = 4 replicates per treatment; mean ± 95 % CI). Rows: ancestral vs. resistance primed community background. Columns: all degraders present; *C. testosteroni* dropped; *P. chlororaphis* dropped; *A. caviae* dropped; or all three degraders dropped. Shaded panels indicate conditions where resistance primed *A. caviae* with accelerated degradation is present. **(C)** Change in the non-*A. caviae* fraction during the main pulse (Δ = day 16 − day 12) in the 20-day experiment. Box plots summarise distributions across replicates (n = 4) by resistance priming background and pre-pulse level (fill colour). Positive values indicate net increase in non-*A. caviae* relative abundance. The centre line is the median, box bounds are the first and third quartiles, and whiskers extend to 1.5× the interquartile range. **Alt text:** Three-panel figure. Panel A shows time-series curves of ampicillin concentration for four species in monoculture; ancestral *A. caviae* degrades more slowly than primed *A. caviae*. Panel B shows a grid of dual-axis plots with *A. caviae* abundance and ampicillin concentration over 24 hours across leave-one-out conditions; ampicillin declines faster in primed than ancestral communities when *A. caviae* is present. Panel C shows box plots of change in non-*A. caviae* fraction during the main pulse, showing positive values (non-*A. caviae* taxa increase) in full-primed communities but negative values (decline) in ancestral communities.

To determine how degrader identity and resistance priming shaped community-level detoxification, we performed a 24 h leave-one-out experiment removing individual or all degraders from ancestral or full-primed community backgrounds (Fig. 5B; Supplementary Fig. S5). Whenever *A. caviae* was present, ampicillin declined significantly earlier in resistance primed than ancestral communities (log-rank *χ^2^* = 23.00, *p* < 0.001; ΔRMST = −12 h). Ancestral communities took the full 24 h culture cycle to degrade ampicillin, and *A. caviae* relative abundance increased throughout this period. In full-primed communities, after degradation (12 h), *A. caviae* relative abundance declined and the non-*A. caviae* fraction increased, indicating that accelerated detoxification transiently released competitors from antibiotic stress even though *A. caviae* maintained dominance.

In line with these results, in the 20-day community experiment, non-*A. caviae* taxa declined during the main pulse in ancestral communities but were maintained or increased in full-primed communities (resistance priming effect: *F*3,48 = 179.2, *p* < 0.001; Supplementary Table S11; Fig. 5C). Increasing pre-pulse priming further increased the non-*A. caviae* fraction across backgrounds (*F*3,48 = 7.67, *p* < 0.001), consistent with prior ecological enrichment of resistant taxa. Together, these results show that resistance priming of the dominant degrader *A. caviae* accelerates environmental detoxification and transiently protects non-degraders, while *A. caviae* itself maintains dominance owing to its high resistance, efficient degradation, and absence of a fitness cost of resistance.

### Mutation and gene expression patterns reflect relaxed and altered selection with resistance priming

Population metagenomic sequencing at the end of the 20-day community experiment revealed that both community-level pre-pulse priming and species-level resistance priming shifted *de novo* mutation profiles (PERMANOVA: pre-pulse *F*1,63 = 2.13, *R^2^*= 0.029, p = 0.023; resistance priming *F*3,63 = 3.65, *R^2^*= 0.15, *p* < 0.001; interaction *F*3,63 = 1.57, *R^2^*= 0.064, *p* = 0.030; Supplementary Tables S12–S13; Fig. 6A,B). The effect was dominated by species-level resistance priming: full-primed communities differed significantly from all others in mutation profiles (all post-hoc *p* < 0.001), while other backgrounds were mutually similar (all post-hoc *p* > 0.30).

**Figure 6.**
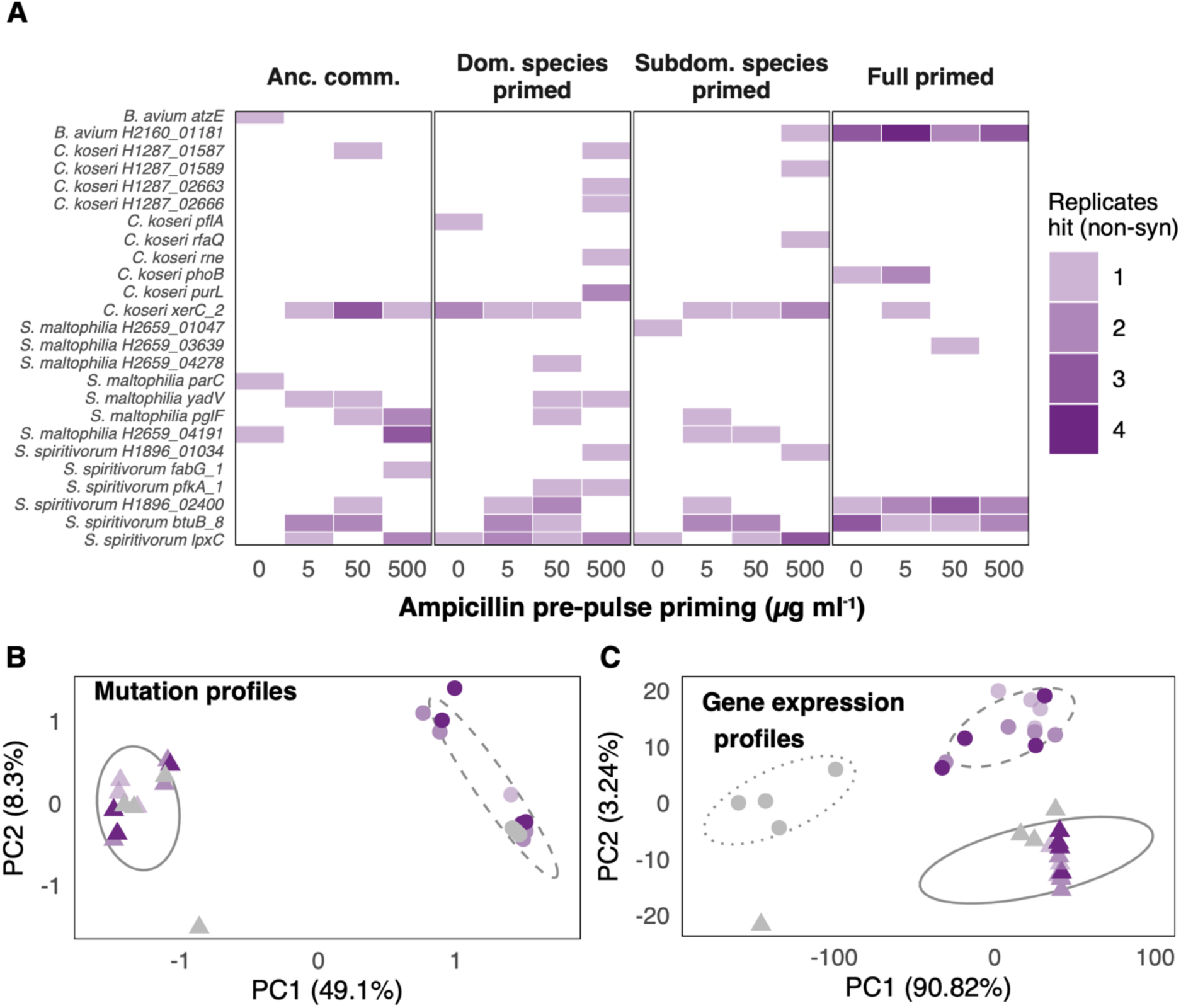
Effect of priming on *de novo* mutations and gene expression. **(A)** Heatmap of high-confidence, recurrent (present in more than one independent community) nonsynonymous *de novo* mutations detected at experimental end-point, across pre-pulse levels (*x*-axis) and resistance priming backgrounds (columns). Tile shade indicates the number of independent replicate communities (1–4) in which the mutation was observed; gene labels include species names. **(B)** Principal component analysis (PCA) of mutation profiles for ancestral and full-primed communities (n = 16 each), with 68 % confidence ellipses shown for each group. **(C)** PCA of gene expression profiles after main ampicillin pulse, normalised by species relative abundances (n = 16 per group), with 68 % confidence ellipses for ancestral and full-primed communities at 5–500 µg mL^−1^ pre-pulse, and ancestral communities at 0 µg mL^−1^ pre-pulse. **Alt text:** Three-panel figure. Panel A shows a heatmap grid of mutation targets (rows) across community types and pre-pulse levels (columns), with darker tiles indicating more replicates carrying each mutation; fully primed communities show fewer mutation targets overall. Panel B shows a PCA scatter plot where ancestral and full-primed communities form separated ellipses, indicating distinct mutation profiles. Panel C shows a PCA scatter plot of gene expression profiles where ancestral and full-primed communities form separated ellipses, with ancestral communities at zero pre-pulse clustering distinctly from antibiotic-exposed ancestral communities.

These differences reflected shifts in mutation targets rather than increased variance (dispersion analysis *F*3,60 = 1.09, *p* = 0.34). For example, in *Citrobacter koseri*, mutations recurred at the *xerC* locus (site-specific recombinase) in ancestral and single-species-primed communities but at *phoB* in full-primed communities; in *Sphingobacterium spiritivorum*, *btuB* was the only recurrent target in full-primed communities but co-occurred with *lpxC* in other backgrounds. Full-primed communities had on average half the number of mutation targets compared with other backgrounds, with more recurrent mutations at the same loci. Similarly, Gene duplications (e.g., *infC* in *Comamonas testosteroni*; *xerC* in *C. koseri* and *Bordetella avium*) occurred predominantly in non-full-primed communities (Supplementary results; Supplementary Fig. S6). These results are consistent with reduced selection pressure arising from prior resistance mutations and narrowing of future evolutionary pathways. Moreover, many targets differed from those observed during monoculture resistance priming, in line with prior work showing that the community context fundamentally alters mutation targets under positive selection [20, 47–50].

Gene expression profiles (RNA-seq) after the main antibiotic pulse, normalised by species relative abundances, differed significantly between ancestral and full-primed communities (PC1: *R* = 0.57, *p* < 0.001, FDR = 0.001; PC2: *R* = −0.71, *p* < 0.001, FDR < 0.001; Fig. 6C), reflecting a distinct community transcriptional state associated with prior resistance evolution. Within ancestral communities, pre-pulse priming had a marginally significant transcriptional effect primarily driven by an antibiotic-free versus antibiotic-exposed contrast (pre-pulse level: PC2 *R* = 0.50, *p* = 0.047, FDR = 0.096). Differentially expressed genes were distributed across treatments without clustering into specific functional pathways, consistent with broad transcriptional adjustments driven by global gene expression changes linked to resistance mutations [51–53] rather than targeted pathway-level responses.

## Discussion

Disturbance responses in microbial communities are frequently interpreted through ecological species sorting or evolutionary change alone [4, 11, 12, 21]. Our results show that in systems where antibiotic degradation transforms the stress environment, these processes are insufficient to explain community dynamics without considering environmental transformation. By separating community-level pre-pulse priming from species-level resistance priming in a unified experimental design, we demonstrate that the two forms of priming act through distinct mechanisms, interact predictably, and together generate a trade-off between resistance and recovery that is shaped, and in unexpected ways modified, by environmental change through antibiotic degradation (Fig. 7).

**Figure 7.**
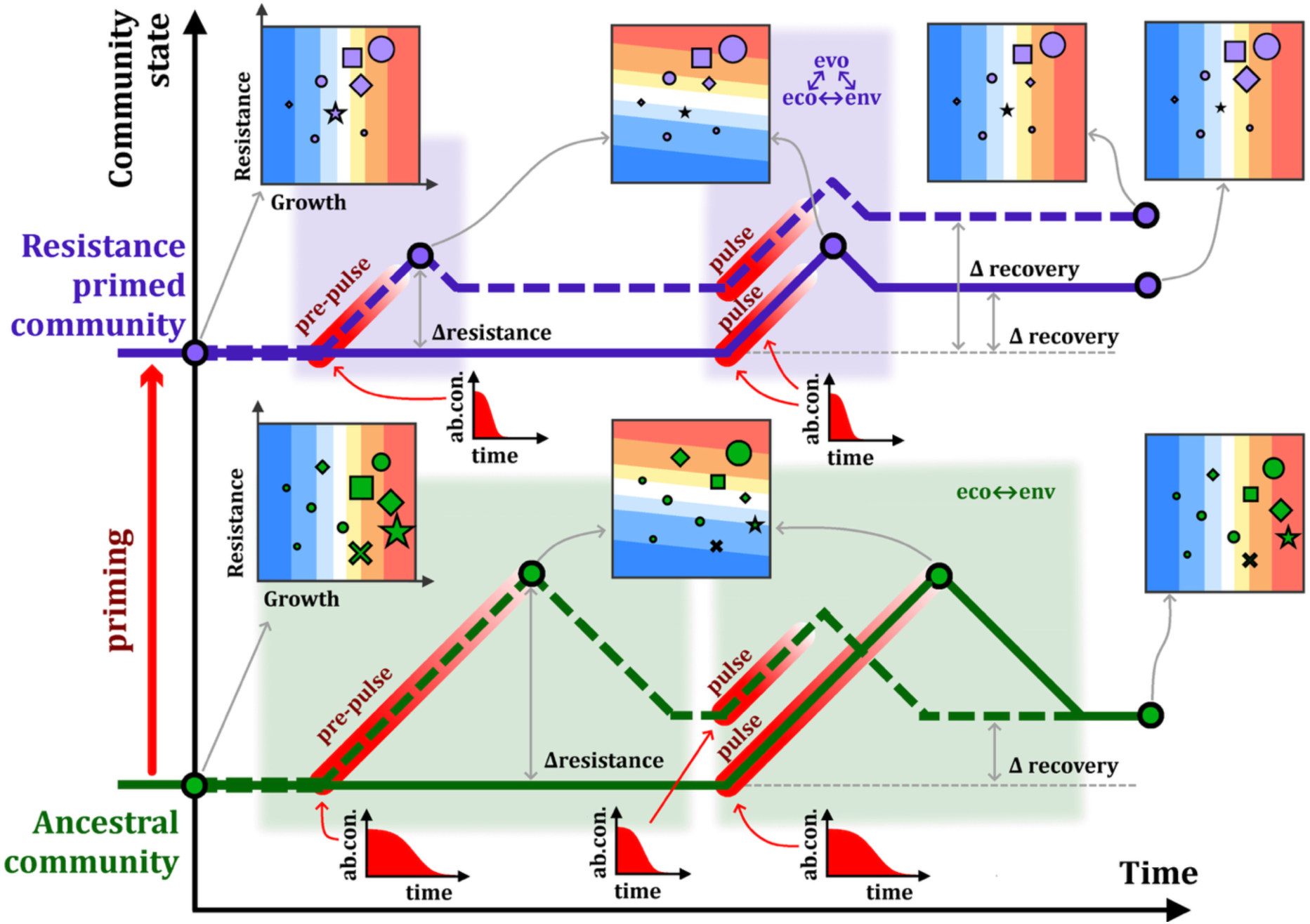
Eco-evolutionary-environmental feedbacks determine resistance and recovery dynamics in an antibiotic degrading community. Species closer to the red zone are favoured. This zone shifts from growth-favouring (no antibiotics) to resistance-favouring (with antibiotics). The resistance primed community is derived from ancestral members (arrow), resulting in a smaller occupied trait space owing to resistance evolution and growth costs. Ancestral community (bottom): Broad trait variation leads to different species being favoured across environments. This drives strong species sorting during pulses, but composition and diversity largely recover after disturbance. Resistance primed community (top): Resistance evolution and faster antibiotic degradation buffer compositional change during pulses. Repeated pulses enrich resistant, competitive taxa, reducing diversity and limiting recovery. **Alt text:** Conceptual diagram with two community scenarios (ancestral, bottom; resistance primed, top) depicted as scatter plots of trait space (growth rate versus resistance). The red selection zone shifts between conditions. Arrows and text describe eco-evo-environmental feedback loops: in the ancestral community, broad trait variation allows recovery; in the resistance primed community, narrowed trait space and faster antibiotic degradation buffer disturbance but limit diversity recovery.

Consistent with ecological filtering predictions [4, 11, 12], community-level pre-pulse priming accelerated compositional sorting toward resistance-enriched states and reduced the magnitude of change required during the main pulse. The observed effects are more parsimoniously explained by trait-based species sorting than by rapid *de novo* resistance evolution, extending our earlier finding on the primacy of species-trait sorting over *de novo* evolution in community responses [21] to a two-pulse scenario and aligning with evidence for constrained evolutionary dynamics in community contexts [47, 54]. Pre-pulse priming therefore modified trajectory speed rather than trajectory outcome, effectively advancing in time the sorting that would ultimately occur during the main pulse regardless. This interpretation is supported by the gene expression data: within ancestral communities, pre-pulse priming produced only a weak transcriptional effect distinguishing antibiotic-free from antibiotic-exposed conditions, offering no evidence for broad evolutionary adaptation prior to the main pulse.

In contrast, species-level resistance priming directly altered the evolutionary state of community members before community assembly. When all species were primed, communities showed markedly reduced compositional fluctuation during antibiotic exposure, consistent with evolutionary buffering [11, 12, 55]. Yet this buffering came at a recovery cost: Shannon diversity declined and dominance structures were reinforced, revealing an ecological cost of evolutionary resistance. Strikingly, ancestral communities recovered as well as or better than full-primed communities despite experiencing stronger compositional change during the pulse. This unexpected result reflects the role of antibiotic degradation in limiting diversity loss in ancestral communities to reversible species sorting rather than local extinction, while the coupling of resistance with competitive dominance in primed communities constrained diversity recovery. The two priming modes thus differentially reshaped the balance between resistance and recovery: pre-pulse priming influenced trajectory dynamics while leaving the final recovered state largely unchanged, whereas resistance priming altered both the disturbance response and the recovery endpoint.

Crucially, neither ecological nor evolutionary priming alone fully explains the observed dynamics. Antibiotic degradation introduced a third axis of community change: environmental transformation. Pre-pulse priming modestly accelerated degradation through enrichment of degraders, an ecological mechanism consistent with compositional carryover. More strikingly, resistance priming of the dominant degrader *A. caviae* accelerated in-community antibiotic breakdown, producing earlier reductions in effective exposure. A potential cause for the accelerated degradation and growth of resistance-primed *A. caviae* is the *ftsI* mutation (penicillin-binding protein 3, PBP3) detected after resistance priming. PBP3 mutations are known to alter β-lactam susceptibility and cell division dynamics [56, 57], potentially increasing growth under antibiotic stress and enabling populations to reach effective degradation thresholds earlier. Consistently, communities containing primed *A. caviae* reached substantially higher cell densities (Supplementary Fig. S2), supporting altered growth dynamics under ampicillin.

This sequence reveals an emergent eco-evolutionary-environmental feedback (Fig. 7). Evolution in a dominant degrader altered environmental dynamics (faster antibiotic removal) [17, 18, 58], which restructured ecological competition (transient protection and expansion of other taxa) [10, 15], which in turn modified selective pressures on the whole community. More broadly, these results demonstrate that microbial disturbance responses are path-dependent not only because of altered trait distributions, but because prior evolutionary history can modify the stress environment itself. In antibiotic-exposed microbiomes and other disturbed systems [1, 22, 59–61], such environmentally mediated evolutionary effects may determine whether communities stabilise, collapse, or reorganise following recurrent stress. The resistance-recovery trade-off identified here is not apparent when examining resistance or recovery in isolation, and may be amplified in natural communities where antibiotic degradation is a common mechanism of community-level tolerance [13, 15]. Future work should examine whether this trade-off holds across different degradation efficiencies, antibiotic classes, and community diversities, and whether eco-evolutionary-environmental feedbacks of this type can be leveraged to steer community composition in applied contexts such as microbiome management or bioremediation.

By integrating ecological filtering, evolutionary priming, and environmental transformation within a unified experimental design, we show that microbial disturbance response cannot be fully understood without considering their interactions. Disturbance history does not merely change who is present; it reshapes how future selection unfolds and how the environment is transformed upon re-exposure. This eco-evolutionary-environmental framework provides a mechanistic basis for predicting how microbial communities respond to recurrent stress and highlights how evolutionary contingency, shaped by prior stress history, determines both resistance and recovery trajectories.

### Data availability

The 16S rRNA gene amplicon sequencing data from the main microbial community evolution experiment are available in the European Nucleotide Archive under accession PRJEB108471, and whole-genome, metagenome and RNA-seq data under the accessions PRJEB109096, PRJEB109023 and PRJEB109068, respectively. Pre-processed data for all downstream analyses is available in GitHub: https://github.com/johannescairns/eco-evo-detox-ampicillin.

### Code availability

The code needed to reproduce the downstream analyses for RNA-seq data is available at https://version.helsinki.fi/biodata-analytics-unit/ampicillin-pulse-experiment.git. The code for reproducing all other downstream analyses and figures is available in GitHub: https://github.com/johannescairns/eco-evo-detox-ampicillin.

## Acknowledgements

We thank Inga-Katariina Aapalampi, Ida Aarni, Emmi Hirvonen, Fanny Koskela, Trine Link, Linda Nevala, Hilla Pellikka, Julia Saloranta, Amanda Silvennoinen, and Milla Similä for technical assistance, and Mikhail Shubin for assistance with drawing figure 7. We wish to acknowledge the Center of Evolutionary applications, University of Turku, Finland, for support in DNA and RNA wet lab methodology; the services of the HiLIFE Biodata Unit supported by HiLIFE and the Biocenter Finland Bioinformatics platform; and CSC – IT Center for Science, Finland, for computational resources. The mass spectrometric analysis was performed at the Turku Metabolomics Centre with the support of Biocenter Finland. This work was supported by the Research Council of Finland (Multidisciplinary Center of Excellence in Antimicrobial Resistance Research, grant #346126; grants #330886 & #327741 to TH; grants #346128 and #364234 to VM), and the SciLifeLab & Wallenberg Data Driven Life Science Program (grant number KAW 2024.0159 to JC).

## Author Contributions Statement

Conceptualization, J.C., T.H., V.M., M.T., V.P.F., and L.B.; Methodology, J.C. and V.M.; Investigation, S.P., O.P., and M.L.; Formal Analysis, J.C., N.S., and R.D.R.; Data Curation, J.C. and N.S.; Writing – Original Draft, J.C.; Writing – Review & Editing, J.C., T.H., V.M., V.P.F., and L.B.; Supervision, J.C., T.H., V.M., M.T., and V.P.F..; Project Administration, J.C. and T.H.; Funding Acquisition, V.M. and T.H.

## References

1. Allison SD, Martiny JBH. Resistance, resilience, and redundancy in microbial communities. Proc Natl Acad Sci USA 2008;105:11512–9. 10.1073/pnas.0801925105

2. Widder S, Allen RJ, Pfeiffer T, Curtis TP, Wiuf C, Sloan WT et al. Challenges in microbial ecology: building predictive understanding of community function and dynamics. ISME J 2016;10:2557–68. 10.1038/ismej.2016.45

3. Lozupone CA, Stombaugh JI, Gordon JI, Jansson JK, Knight R. Diversity, stability and resilience of the human gut microbiota. Nature 2012;489:220–30. 10.1038/nature11550

4. Thorogood R, Mustonen V, Aleixo A, Aphalo PJ, Asiegbu FO, Cabeza M et al. Understanding and applying biological resilience, from genes to ecosystems. npj Biodivers 2023;2:8. 10.1038/s44185-023-00022-6

5. Becks L, Ellner SP, Jones LE, Hairston NG. The functional genomics of an eco-evolutionary feedback loop: linking gene expression, trait evolution, and community dynamics. Ecol Lett 2012;15:492–501. 10.1111/j.1461-0248.2012.01763.x

6. Yoshida T, Jones LE, Ellner SP, Fussmann GF, Hairston NG. Rapid evolution drives ecological dynamics in a predator-prey system. Nature 2003;424:303–6. 10.1038/nature01767

7. Marsland R, Cui W, Goldford J, Sanchez A, Korolev K, Mehta P. Available energy fluxes drive a transition in the diversity, stability, and functional structure of microbial communities. PLoS Comput Biol 2019;15:e1006793. 10.1371/journal.pcbi.1006793

8. Wittebolle L, Marzorati M, Clement L, Balloi A, Daffonchio D, Heylen K et al. Initial community evenness favours functionality under selective stress. Nature 2009;458:623–6. 10.1038/nature07840

9. Beyter D, Tang PZ, Becker S, Hoang T, Bilgin D, Lim YW et al. Diversity, productivity, and stability of an industrial microbial ecosystem. Appl Environ Microbiol 2016;82:2494–505. 10.1128/AEM.03965-15

10. Denk-Lobnig M, Wood KB. Antibiotic resistance in bacterial communities. Curr Opin Microbiol 2023;74:102306. 10.1016/j.mib.2023.102306

11. Xu CCY, Fugère V, Barbosa da Costa N, Beisner BE, Bell G, Cristescu ME et al. Pre-exposure to stress reduces loss of community and genetic diversity following severe environmental disturbance. Curr Biol 2025;35:1061–73.e4. 10.1016/j.cub.2025.01.037

12. Cairns J, Hogle S, Alitupa E, Mustonen V, Hiltunen T. Pre-exposure of abundant species to disturbance improves resilience in microbial metacommunities. Nat Ecol Evol 2025;9:395–405. 10.1038/s41559-024-02624-0

13. Ma HR, Xu HZ, Kim K, Anderson DJ, You L. Private benefit of β-lactamase dictates selection dynamics of combination antibiotic treatment. Nat Commun 2024;15:8799. 10.1038/s41467-024-52711-w

14. Cairns J, Koskinen K, Penttinen R, Patinen T, Hartikainen A, Jokela R et al. Black Queen evolution and trophic interactions determine plasmid survival after the disruption of the conjugation network. mSystems 2018;3:e00104–18. 10.1128/msystems.00104-18

15. Pathak A, Angst DC, León-Sampedro R, Hall AR. Antibiotic-degrading resistance changes bacterial community structure via species-specific responses. ISME J 2023;17:1495–503. 10.1038/s41396-023-01465-2

16. Wetherington MT, Copeland R, Zhang C, Hammer BK, Yunker PJ. Higher levels of antibiotic resistance are less competitive: the hidden ecological cost of no-metabolic cost resistance. bioRxiv 2025, 2025.12.15.694367. 10.64898/2025.12.15.694367

17. Fröhlich C, Chen JZ, Gholipour S, Erdogan AN, Tokuriki N. Evolution of β-lactamases and enzyme promiscuity. Protein Eng Des Sel 2021;34:gzab013. 10.1093/protein/gzab013

18. Petrosino J, Cantu C, Palzkill T. β-Lactamases: protein evolution in real time. Trends Microbiol 1998;6:323–7. 10.1016/S0966-842X(98)01317-1

19. Quinn AM, Bottery MJ, Thompson H, Friman VP. Resistance evolution can disrupt antibiotic exposure protection through competitive exclusion of the protective species. ISME J 2022;16:2433–47. 10.1038/s41396-022-01285-w

20. Muzafar S, Nair RR, Andersson DI, Warsi OM. Interspecies interaction alters the trajectory of antibiotic resistance evolution by amplifying negative fitness epistasis. ISME J 2026;20:wrag014. 10.1093/ismejo/wrag014

21. Cairns J, Jokela R, Becks L, Mustonen V, Hiltunen T. Repeatable ecological dynamics govern the response of experimental communities to antibiotic pulse perturbation. Nat Ecol Evol 2020;4:1385–94. 10.1038/s41559-020-1272-9

22. Kivikoski M, Cairns J, Hogle SL, Pausio S, Becks L, Mustonen V, et al. Evolution induced state shifts in a long-term microbial community experiment. Proc Natl Acad Sci USA 2026;123:e2533269123. 10.1073/pnas.2533269123

23. Good BH, Rosenfeld LB. Eco-evolutionary feedbacks in the human gut microbiome. Nat Commun 2023;14:7146. 10.1038/S41467-023-42769-3

24. Dethlefsen L, Relman DA. Incomplete recovery and individualized responses of the human distal gut microbiota to repeated antibiotic perturbation. Proc Natl Acad Sci USA 2011;108:4554–61. 10.1073/pnas.1000087107

25. Hogle SL, Ruusulehto L, Cairns J, Hultman J, Hiltunen T. Localized coevolution between microbial predator and prey alters community-wide gene expression and ecosystem function. ISME J 2023;17:514–24. 10.1038/s41396-023-01361-9

26. Cairns J, Jokela R, Hultman J, Tamminen M, Virta M, Hiltunen T. Construction and characterization of synthetic bacterial community for experimental ecology and evolution. Front Genet 2018;9:312. 10.3389/fgene.2018.00312

27. Hogle SL, Tamminen M, Hiltunen T. Complete genome sequences of 30 bacterial species from a synthetic community. Microbiol Resour Announc 2024;13:e00111–24. 10.1128/mra.00111-24

28. Glenn TC, Pierson TW, Bayona-Vásquez NJ, Kieran TJ, Hoffberg SL, Thomas JC et al. Adapterama II: universal amplicon sequencing on Illumina platforms (TaggiMatrix). PeerJ 2019;7:e7786. 10.7717/peerj.7786

29. Bischofberger AM, Cairns J, Aapalampi IK, Pausio S, Lindqvist M, Mustonen V et al. Community state shifts driven by total carbon availability over resource complexity in a synthetic microbial community. ISME Commun 2026, ycag149, 10.1093/ismeco/ycag149

30. Hogle SL, Hepolehto I, Ruokolainen L, Cairns J, Hiltunen T. Effects of phenotypic variation on consumer coexistence and prey community structure. Ecol Lett 2022;25:307–19. 10.1111/ele.13924

31. Li H. Aligning sequence reads, clone sequences and assembly contigs with BWA-MEM. arXiv 2013, 1303.3997. 10.48550/arXiv.1303.3997

32. Benjamin D, Sato T, Cibulskis K, Getz G, Stewart C, Lichtenstein L. Calling somatic SNVs and indels with Mutect2. bioRxiv 2019, 861054. 10.1101/861054

33. Cingolani P, Platts A, Coon M, Nguyen T, Wang L, Land SJ et al. A program for annotating and predicting the effects of single nucleotide polymorphisms, SnpEff: SNPs in the genome of Drosophila melanogaster strain w1118; iso-2; iso-3. Fly (Austin) 2012;6:80–92. 10.4161/fly.19695

34. Jugas R, Sedlar K, Vitek M, Nykrynova M, Barton V, Bezdicek M et al. CNproScan: hybrid CNV detection for bacterial genomes. Genomics 2021;113:3103–11. 10.1016/j.ygeno.2021.06.040

35. Patro R, Duggal G, Love MI, Irizarry RA, Kingsford C. Salmon provides fast and bias-aware quantification of transcript expression. Nat Methods 2017;14:417–9. 10.1038/nmeth.4197

36. Patel H, Ewels P, Peltzer A, Manning J, Botvinnik O, Sturm G et al. nf-core/rnaseq: nf-core/rnaseq v3.14.0 - Hassium Honey Badger. Zenodo 2024. 10.5281/zenodo.10471647

37. Love MI, Huber W, Anders S. Moderated estimation of fold change and dispersion for RNA-seq data with DESeq2. Genome Biol 2014;15:550. 10.1186/S13059-014-0550-8

38. R Core Team. R: A Language and Environment for Statistical Computing. R Foundation for Statistical Computing, 2024.

39. Oksanen J, Simpson GL, Blanchet FG, Kindt R, Legendre P, Minchin PR et al. Community Ecology Package. R package version 2.7–1, 2025. https://vegandevs.github.io/vegan/

40. Pinheiro J, Bates D, DebRoy S, Sarkar D, R Core Team. nlme: Linear and Nonlinear Mixed Effects Models. R package, 2021. https://CRAN.R-project.org/package=nlme

41. Lenth RV, Piaskowski J. emmeans: Estimated Marginal Means, aka Least-Squares Means. R package, 2017. https://github.com/rvlenth/emmeans

42. Venables WN, Ripley BD. Modern Applied Statistics with S, 4th ed. Springer, 2002.

43. Krijthe JH. Rtsne: T-Distributed Stochastic Neighbor Embedding using a Barnes-Hut Implementation. R package, 2015. https://github.com/jkrijthe/Rtsne

44. van der Maaten L. Accelerating t-SNE using tree-based algorithms. J Mach Learn Res 2014;15:3221–45.

45. Pinheiro F, Warsi O, Andersson DI, Lässig M. Metabolic fitness landscapes predict the evolution of antibiotic resistance. Nat Ecol Evol 2021;5:677–87. 10.1038/s41559-021-01397-0

46. Melnikov SV, Stevens DL, Fu X, Kwok HS, Zhang JT, Shen Y et al. Exploiting evolutionary trade-offs for posttreatment management of drug-resistant populations. Proc Natl Acad Sci USA 2020;117:17924–31. 10.1073/pnas.2003132117

47. Scheuerl T, Hopkins M, Nowell RW, Rivett DW, Barraclough TG, Bell T. Bacterial adaptation is constrained in complex communities. Bacterial adaptation is constrained in complex communities. Nat Commun 2020;11:754. 10.1038/s41467-020-14570-z

48. Leónidas Cardoso L, Durão P, Amicone M, Gordo I. Dysbiosis individualizes the fitness effect of antibiotic resistance in the mammalian gut. Nat Ecol Evol 2020;4:1268–78. 10.1038/s41559-020-1235-1

49. Fang P, Elena AX, Kunath MA, Berendonk TU, Klümper U. Reduced selection for antibiotic resistance in community context is maintained despite pressure by additional antibiotics. ISME Commun 2023;3:52. 10.1038/s43705-023-00262-4

50. Klümper U, Recker M, Zhang L, Yin X, Zhang T, Buckling A et al. Selection for antimicrobial resistance is reduced when embedded in a natural microbial community. ISME J 2019;13:2927–37. 10.1038/s41396-019-0483-z

51. Sun L, Ashcroft P, Ackermann M, Bonhoeffer S. Stochastic gene expression influences the selection of antibiotic resistance mutations. Mol Biol Evol 2020;37:58–70. 10.1093/molbev/msz199

52. Trauner A, Banaei-Esfahani A, Gygli SM, Warmer P, Feldmann J, Zampieri M et al. Expression dysregulation as a mediator of fitness costs in antibiotic resistance. Antimicrob Agents Chemother 2021;65:e00504–21. 10.1128/AAC.00504-21

53. Suzuki S, Horinouchi T, Furusawa C. Prediction of antibiotic resistance by gene expression profiles. Nat Commun 2014;5:5792. 10.1038/ncomms6792

54. Bottery MJ, Pitchford JW, Friman VP. Ecology and evolution of antimicrobial resistance in bacterial communities. ISME J 2021;15:939–48. 10.1038/s41396-020-00832-7

55. Hernández-Navarro L, Asker M, Rucklidge AM, Mobilia M. Coupled environmental and demographic fluctuations shape the evolution of cooperative antimicrobial resistance. J R Soc Interface 2023;20:20230393. 10.1098/rsif.2023.0393

56. Deatherage BL, Lara JC, Bergsbaken T, Barrett SLR, Lara S, Cookson BT. Biogenesis of bacterial membrane vesicles. Mol Microbiol 2009;72:1395–407. 10.1111/j.1365-2958.2009.06731.x

57. Kim SW, Park SB, Im SP, Lee JS, Jung JW, Gong TW et al. Outer membrane vesicles from β-lactam-resistant *Escherichia coli* enable the survival of β-lactam-susceptible *E. coli* in the presence of β-lactam antibiotics. Sci Rep 2018;8:5402. 10.1038/S41598-018-23656-0

58. Gross R, Yelin I, Lázár V, Datta MS, Kishony R. Beta-lactamase dependent and independent evolutionary paths to high-level ampicillin resistance. Nat Commun 2024;15: 5383. 10.1038/s41467-024-49621-2

59. Flemming HC, Wuertz S. Bacteria and archaea on Earth and their abundance in biofilms. Nat Rev Microbiol 2019;17:247–60. 10.1038/s41579-019-0158-9

60. Prosser JI, Bohannan BJM, Curtis TP, Ellis RJ, Firestone MK, Freckleton RP et al. The role of ecological theory in microbial ecology. Nat Rev Microbiol 2007;5:384–92. 10.1038/nrmicro1643

61. Shade A, Peter H, Allison SD, Baho DL, Berga M, Bürgmann H et al. Fundamentals of microbial community resistance and resilience. Front Microbiol 2012;3:417. 10.3389/fmicb.2012.00417

