## Supplementary Information for "Eco-evolutionary dynamics and environmental detoxification jointly shape bacterial community response to antibiotic perturbation"

This PDF file includes:

- Supplementary Methods:
  - Detailed bioinformatics pipeline parameters
- Supplementary Results:
  - Detailed genomic changes associated with species-level resistance priming
  - Copy number variants
- Supplementary Figures S1–S6
- Supplementary Tables S1–S13

**Supplementary Methods**

Detailed bioinformatics pipeline parameters

*16S rRNA amplicon sequence processing*

Community composition was analysed from 16S rRNA gene amplicon data following a previously established protocol [1]. Briefly, library indices were demultiplexed using Illumina bcl2fastq v2.2. Paired-end reads were quality trimmed, merged, and filtered using standard criteria, then mapped to reference sequences from the 23 community members using a custom pipeline. Species counts were normalised by 16S rRNA gene copy number obtained from genome annotations.

*Whole-genome sequencing (WGS) variant calling*

WGS data were processed on the CSC computing cluster Puhti adapting a previously published pipeline [2]. FASTQ files were pre-processed using fastp v0.23.4 [3] with default settings. Reads were aligned to a custom HAMBI reference genome database [4]. using BWA-MEM v0.7.17 [5]. SAM-to-BAM conversion, read group addition, sorting, and duplicate marking used SAMtools v1.16.1 [6] and GATK v4.4.0 tools SortSam, MarkDuplicates, and BuildBamIndex. Variant calling used GATK Mutect2 [7] with a maximum of 75 reads per alignment start position; variants were filtered using FilterMutectCalls in microbial mode. Variants were annotated using SnpEff v5.1f [8] and converted to tabular format using SnpSift v5.1f [9] and GATK VariantsToTable. Only protein-altering (nonsynonymous) variants were retained for downstream analysis.

*Metagenomic variant calling*

Metagenomic processing followed a similar approach to WGS with the following modifications. After fastp pre-processing [3], reads were partitioned by species using BBMap v39.01 [10]; species with < 5% coverage in a sample were excluded. BWA-MEM alignment was performed against the custom HAMBI reference database, followed by SAMtools and GATK processing as described above. Additional hard filters were applied separately to SNPs and INDELs using GATK

VariantFiltration: for SNPs, filters excluded calls with > 3 SNPs per 35 bases, SOR > 3.0, FS > 60.0, MQ < 40.0, MQRankSum < −12.5, or ReadPosRankSum < −8.0; for INDELs, > 3 INDELs per 35 bases, FS > 200.0, or ReadPosRankSum < −20.0. SNPs or INDELs within 10 bases of an INDEL and SNPs within 3 bases of another variant were removed using BCFtools [6]. Two analysis inputs were derived from the filtered variant set: (i) a presence/absence matrix of unique protein-altering mutation targets per community replicate (for PERMANOVA), and (ii) per-replicate counts of unique mutation targets (for Poisson regression modelling).

*Background and artefact filtering*: putative laboratory-adaptation or common background targets observed in the control community were discarded unless they also appeared in the curated WGS set. Targets were retained only if they showed strong evidence of presence in min. one sample (stringent depth and allele-frequency support) or matched a high-confidence WGS target. Targets with over one independent hit within the same population or observed across over 40 community samples (two-thirds of all communities) were excluded.

*Copy number variant (CNV) analysis*

CNVs were evaluated from WGS and metagenomic BAM files using the CNproScan package v0.999 [11], normalised for GC content and genome mappability. Read depth and mappability files were generated using SAMtools v1.16.1 [6] and GenMap v1.3.0 [12], respectively.

*Filtering:* For whole-genome data (resistance primed monoculture clones), we focused on high-confidence duplications based on read depth (average coverage ≥ 100×), copy number ≥ 2, event length ≥ 800 bp, and duplication support ≥ 2 reads. For metagenomic samples from the community experiment, we focused on recurrent gene duplications detected in ≥ 2 independent community replicates with filtering criteria: copy number ≥ 2, event length ≥ 300 bp, duplication read support ≥ 5, total reads ≥ 20, and ≥ 50 % overlap with an annotated gene.

*RNA-seq processing*

Raw RNA-seq reads were processed and pseudoaligned using the nf-core rnaseq pipeline v3.14.0 [13, 14] with default parameters (rseqc quality control was skipped as it is not recommended for prokaryotes). Pseudoalignment was performed with Salmon v1.10.1 [15] against a custom HAMBI WGS database [4]. A custom normalisation factor based on relative species abundances was incorporated into the gene expression analysis [16]. Principal component analysis of gene expression profiles was performed and principal components were correlated with covariate metadata using DEGreport v1.40.1 [17, 18]. Differential expression analysis used DESeq2 v1.44.0 [19]; genes were considered differentially expressed when the adjusted $p$-value was $< 0.05$ and the shrunken log2 fold change was $> 1$ or $< -1$. Functional annotation used EggNOG-mapper v2.0.0 [20] with EggNOG v5.0 [21], supplemented by the Comprehensive Antibiotic Resistance Database (CARD) v4.0.1 [22]. Functional enrichment of differentially expressed genes used the hypergeometric test in clusterProfiler v4.12.0 [23], summarising KEGG orthology identifiers to KEGG pathways; pathways with FDR-adjusted $p$-values $< 0.05$ were considered significantly enriched.

### Supplementary Results

Detailed genomic changes associated with species-level resistance priming

To characterise the genetic basis of phenotypic changes observed after resistance priming, we whole-genome sequenced resistance primed clones from 10 dominant species (*Aeromonas caviae, Sphingobacterium spiritivorum, Citrobacter koseri, Comamonas testosteroni, Bordetella avium, Morganella morganii, Stenotrophomonas maltophilia, Agrobacterium tumefaciens, Paracoccus denitrificans, and Pseudomonas chlororaphis*). Fixed nonsynonymous mutations or large structural variants were detected in six species.

*Species with increased MIC and increased or stable k*

*Aeromonas caviae* carried mutations in *ftsI* (encoding penicillin-binding protein 3, PBP3) and *erfK*, both involved in peptidoglycan cross-linking and known to confer β-lactam resistance via bypass of penicillin-binding protein activity [24, 25], as well as in *rpoA* (RNA polymerase alpha subunit), linked to β-lactamase-independent high-level resistance [26]. This species was unique in showing increased MIC without a significant *k* cost, suggesting resistance mutations that impose minimal growth trade-offs under antibiotic-free conditions.

*Sphingobacterium spiritivorum* showed a large-scale gene amplification (13× copy number increase) affecting a genomic block containing the β-lactamase gene *blaB* [27] and several flanking genes (*aguA, lpxB, sigV, mpl, nagA, nahK, rpmE*, and *tpiA*). This amplification likely confers resistance through increased β-lactamase production. A corresponding increase in MIC was observed in this species.

*Species with increased MIC and decreased k*

*Bordetella avium* carried a mutation in *cysS* (cysteine-tRNA ligase), previously associated with stress response and reduced antibiotic susceptibility via effects on translational fidelity [28–30].

*Pseudomonas chlororaphis* carried mutations in rne [31] (encoding RNase E, potentially reducing OmpA porin abundance via altered mRNA processing, which could confer resistance via reduced outer membrane permeability [32]) and *ampR* [33] (a LysR-type transcriptional regulator of AmpC β-lactamase production). This species showed one of the strongest resistance-associated *k* costs among abundant community members.

*Species with negligible or minor MIC increase and decreased k*

*Stenotrophomonas maltophilia* harboured a mutation in *smf-1*, implicated in biofilm-associated resistance mechanisms [34] but likely reflecting general laboratory adaptation given its recurrence in this species in a prior study without antibiotic exposure [35] and the negligible MIC change observed here.

*Agrobacterium tumefaciens* carried a mutation in *rbsA*, a component of the ribose ATP-binding cassette transporter system, potentially mediating efflux through ABC transporter-mediated mechanisms [36, 37].

*Species without detectable fixed mutations*

The remaining four sequenced species (*C. koseri, C. testosteroni, M. morganii, and P. denitrificans*) showed trait shifts without fixed nonsynonymous mutations. This could reflect transient genetic changes (e.g., gene amplifications not captured as fixed mutations in our pipeline), large-scale structural rearrangements, or phenotypic responses mediated by inducible mechanisms such as epigenetic modification [38].

Copy number variants

We analysed copy number variants (CNVs) from whole-genome sequencing (WGS) of monoculture resistance primed clones and from population metagenomics of community experiment samples to assess the contribution of gene duplications to antibiotic resistance and adaptation [38].

*WGS (resistance primed monoculture clones)*

One prominent large-scale duplication was detected: the 13× amplification of a genomic block in *S. spiritivorum* containing the β-lactamase gene *blaB* and flanking genes (described in Supplementary results above).

*Metagenomics (community experiment samples)*

Recurrent duplications occurred in four species:

- *Comamonas testosteroni*: duplication of *infC* (translation initiation factor IF-3), previously shown to be upregulated under antibiotic stress [39], occurred predominantly in ancestral and single-species-primed community backgrounds.
- *Citrobacter koseri*: duplication of *xerC* (site-specific recombinase), also a point-mutation target in the community experiment (Fig. 6A), and implicated in antibiotic-associated DNA damage repair [40], occurred in multiple non-full-primed communities.
- *Bordetella avium*: duplication of *xerC*, as in *C. koseri*, detected in ancestral communities.

The predominant occurrence of gene duplications in non-full-primed community backgrounds (Supplementary Fig. S6) is consistent with stronger antibiotic selection in ancestral communities, where more *de novo* adaptive changes are expected. In full-primed communities, pre-existing resistance mutations likely reduced selection pressure, leading to fewer CNV events.

### Supplementary Figures

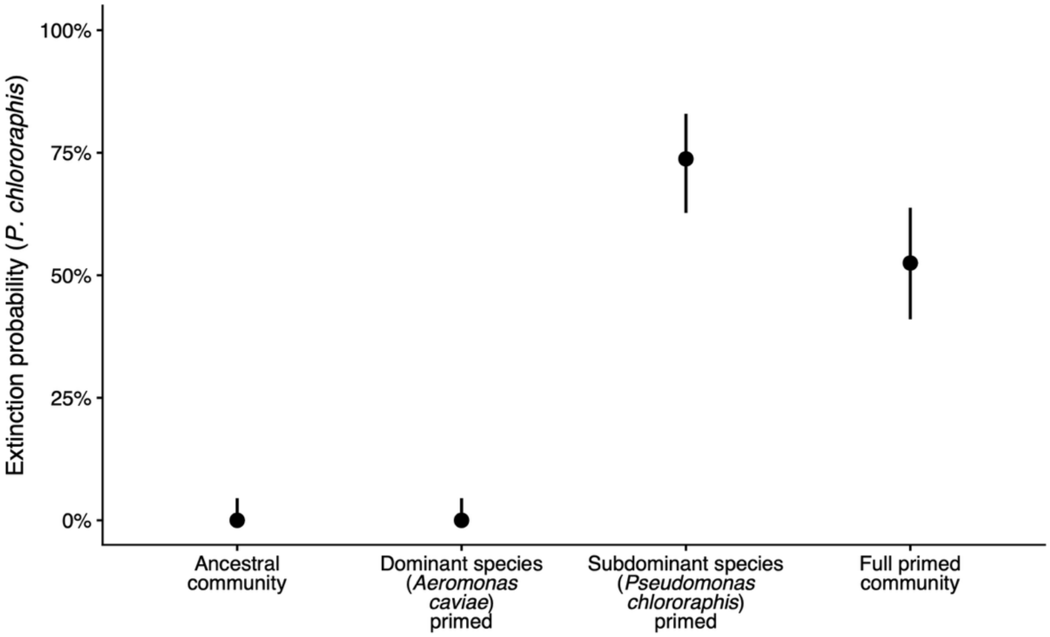


**Figure S1 | Extinction of *Pseudomonas chlororaphis* across species-level resistance priming backgrounds.** Estimated probability that *P. chlororaphis* was extinct during the 20-day serial-passage experiment, shown for each community type varying the resistance priming background of member species. Extinct was defined a priori as zero mapped reads for the focal species in a sample (relative abundance = 0). For each background, extinction probability was computed as the proportion of samples in which the species was extinct, pooling across all pre-pulse ampicillin concentrations (0, 5, 50, and 500 µg $mL^{-1}$) and across all scheduled sampling days (4, 8, 12, 16, and 20). Points show the proportion (k/n) and vertical lines show two-sided 95 % binomial confidence intervals calculated with the Wilson score method. Nominal sample size per background was n = 80 (4 biological replicates × 4 pre-pulse levels × 5 days); any missing or excluded samples were removed from both numerator and denominator. This result complements Fig. 2A in the main text by quantifying how the growth cost of resistance in the resistance primed community member *P. chlororaphis* translated into frequent loss of that species from communities.

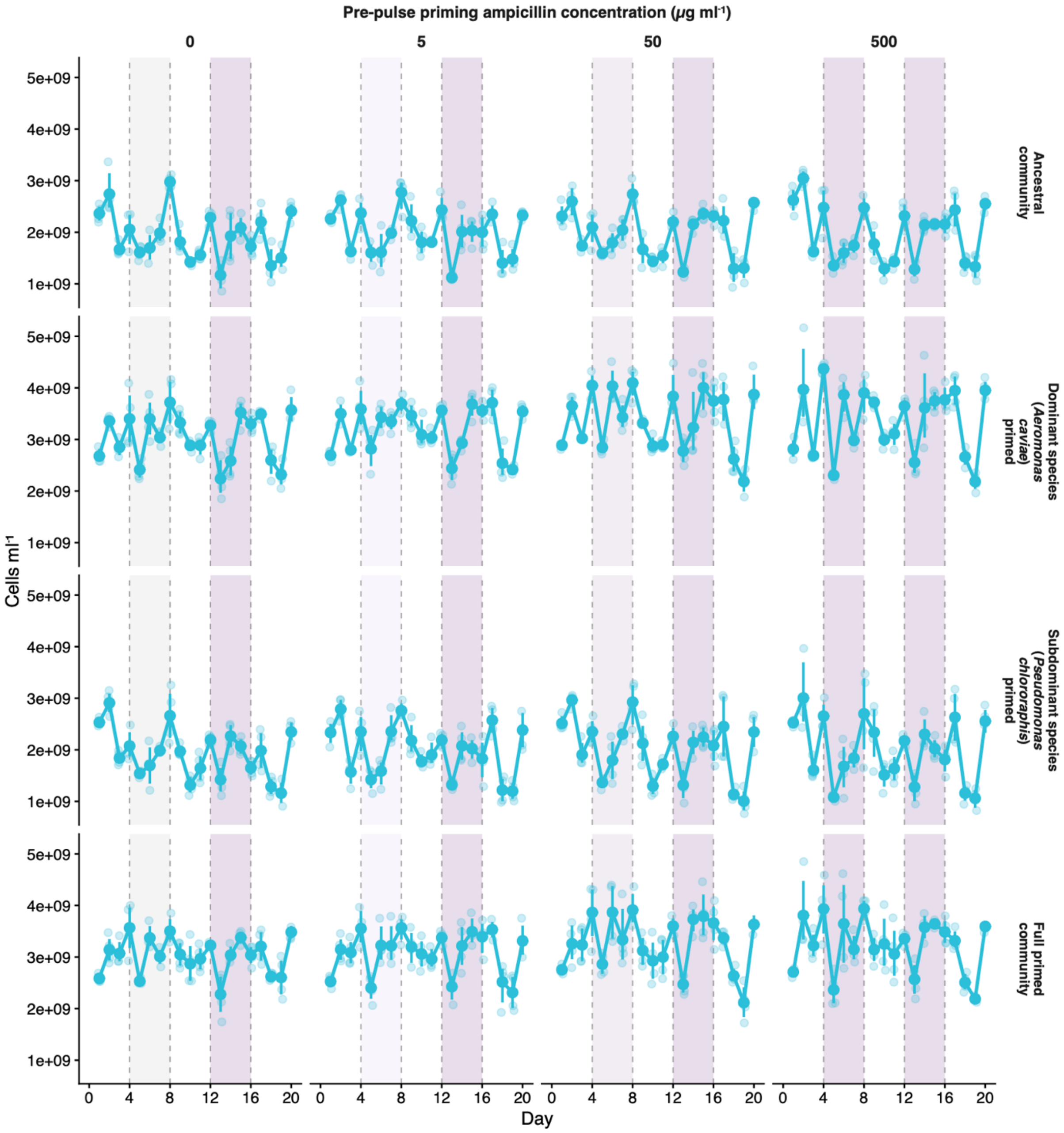


**Figure S2 | Cell density over 20-day ampicillin pulse experiment across community-level pre-pulse priming levels and species-level resistance priming backgrounds.** Cell density (cells mL$^{-1}$) is shown over time (days 0–20) for replicate communities (points; n = 4 per treatment), with the mean across replicates (solid line) and nonparametric bootstrapped 95 % confidence intervals (thin error bars around the mean). Columns indicate the pre-pulse ampicillin concentration (µg mL$^{-1}$): 0, 5, 50, and 500. Rows indicate the community type: ancestral community; dominant species *Aeromonas caviae* resistance primed; subdominant species *Pseudomonas chlororaphis* primed; and full primed community. Experimental phases are marked along the x-axis: acclimation (days 0–4), pre-pulse (days 4–8; deepening purple shading as a function of pre-pulse intensity), intermediate recovery (days 8–12), main pulse (days 12–16; deepest purple shading), and final recovery (days 16–20). Vertical dashed lines at days 4, 8, 12, and 16 indicate phase boundaries.

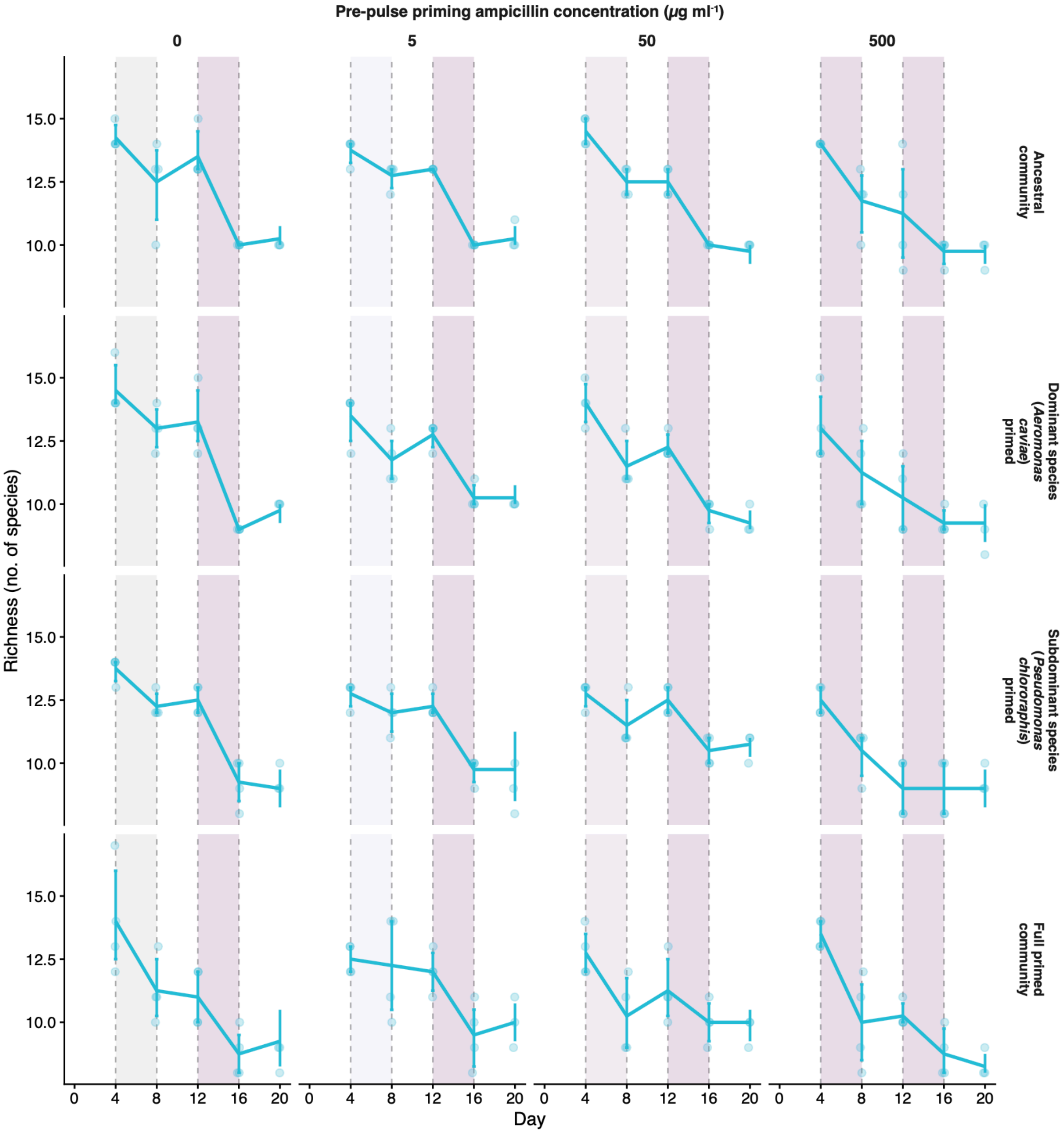


**Figure S3 | Species richness over two-pulse community experiment across community-level pre-pulse priming levels and species-level resistance priming backgrounds.** Points show the number of species detected per community (richness; count of taxa with non-zero relative abundance) at each sampling day (4, 8, 12, 16, 20) across the 20-day serial-passage experiment. The solid black line is the treatment mean and vertical error bars are bootstrapoed 95 % confidence intervals. Each treatment (pre-pulse priming concentration × community type) was propagated in four independent biological replicates (n = 4 per panel; points jittered horizontally for visibility). Columns indicate the pre-pulse ampicillin concentration (μg mL$^{-1}$): 0, 5, 50, and 500. Rows indicate the community background: Ancestral community; dominant species *Aeromonas caviae* resistance primed; subdominant species *Pseudomonas chlororaphis* primed; and full primed community. Experimental phases are marked along the x-axis: acclimation (days 0–4), pre-pulse (days 4–8; deepening purple shading as a function of pre-pulse intensity), intermediate recovery (days 8–12), main pulse (days 12–16; deepest purple shading), and final recovery (days 16–20). Vertical dashed lines at days 4, 8, 12, and 16 indicate phase boundaries.

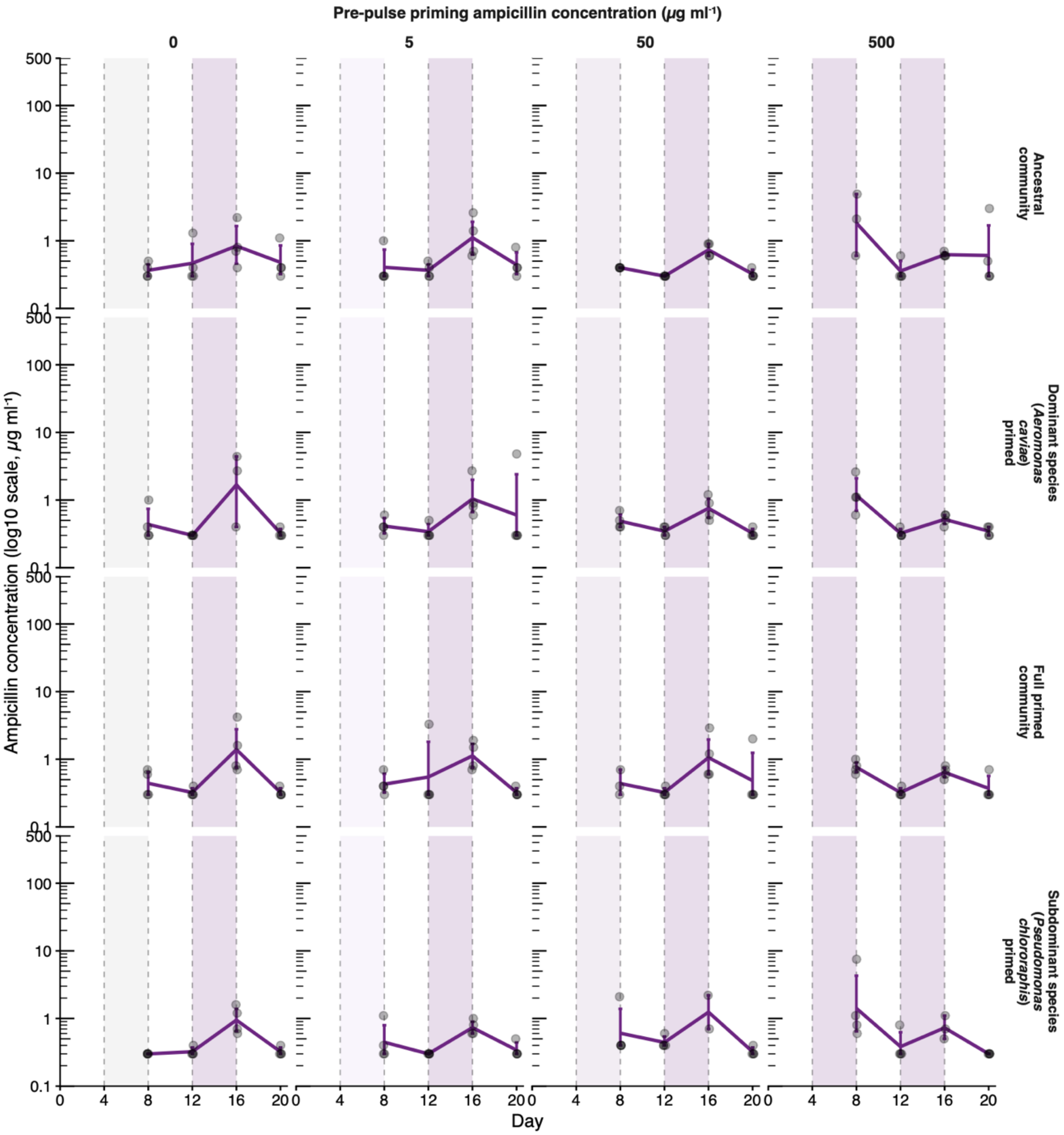


**Figure S4 | Ampicillin dynamics over two-pulse community experiment across community-level pre-pulse priming levels and species-level resistance priming backgrounds.** Ampicillin concentration (μg $mL^{-1}$) is shown over time (days 0–20) for replicate communities (points; n = 4 per treatment), with the mean (solid line) and bootstrapped 95 % confidence intervals. The y-axis is square-root transformed and constrained to 0–500 μg $mL^{-1}$ to show both the maximum pulse magnitude and low-concentration differences. Columns indicate the pre-pulse ampicillin concentration (μg $mL^{-1}$): 0, 5, 50, and 500. Rows indicate the community type: ancestral community; dominant species *Aeromonas caviae* resistance-primed; subdominant species *Pseudomonas chlororaphis* primed; and full primed community. Experimental phases are marked along the x-axis: acclimation (days 0–4), pre-pulse (days 4–8; deepening purple shading as a function of pre-pulse intensity), intermediate recovery (days 8–12), main pulse (days 12–16; deepest purple shading), and final recovery (days 16–20). Vertical dashed lines at days 4, 8, 12, and 16 indicate phase boundaries. The concentrations were measured with LC/MS after a culture cycle and therefore reflect the level to which ampicillin was degraded by the community (bacteria-free controls show no ampicillin decline) from a maximum concentration of 500 μg $mL^{-1}$ (highest pre-pulse and main pulse level).

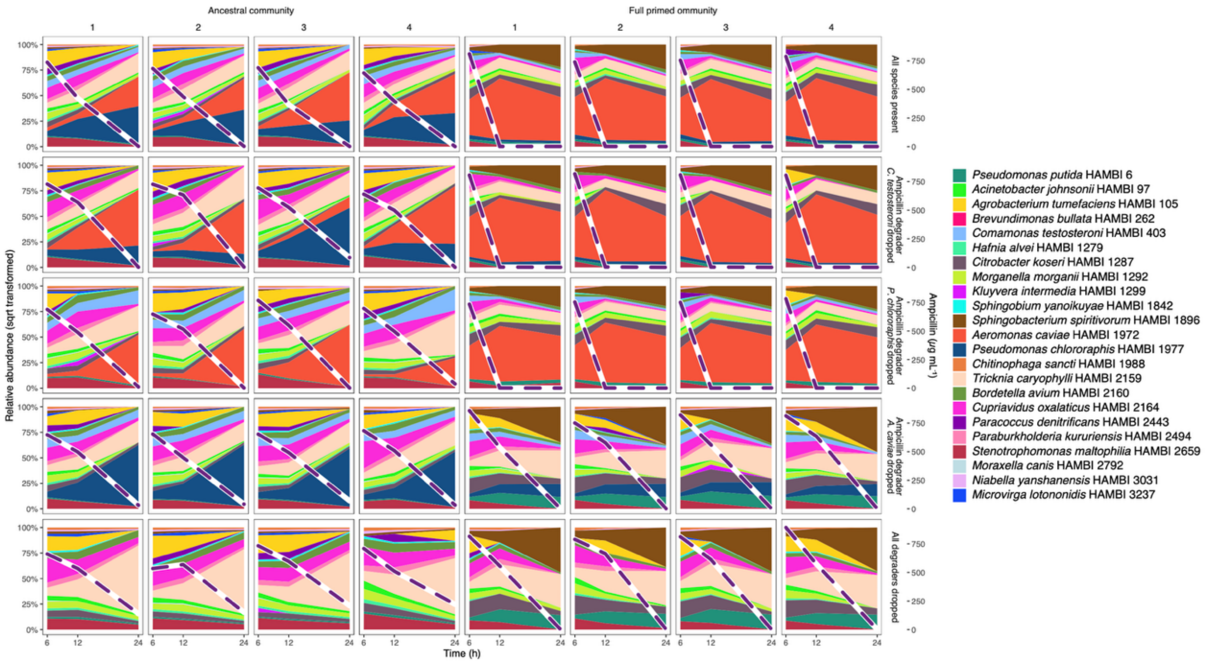


**Figure S5 | Community composition and ampicillin dynamics in 24-h leave-one-out experiment.** Community composition is shown for each replicate as stacked area plots of species relative abundance at 6, 12, and 24 h. Species fractions were square-root transformed and renormalised to sum to 1 within each sample to enhance visibility of low-abundance taxa while preserving relative differences among species. The dashed purple line indicates ampicillin concentration (µg mL$^{-1}$; right y-axis), overlaid on the same panels. Rows indicate degrader-removal treatments: all species present; ampicillin degrader *Comamonas testosteroni* removed; *Pseudomonas chlororaphis* removed; *Aeromonas caviae* removed; and all three degraders removed. Columns indicate community background (ancestral community vs. full primed community) and biological replicate (n = 4 per background × removal treatment). The duration of the leave-one-out experiment (24 h) corresponds to the duration of one culture cycle in two-pulse community experiment.

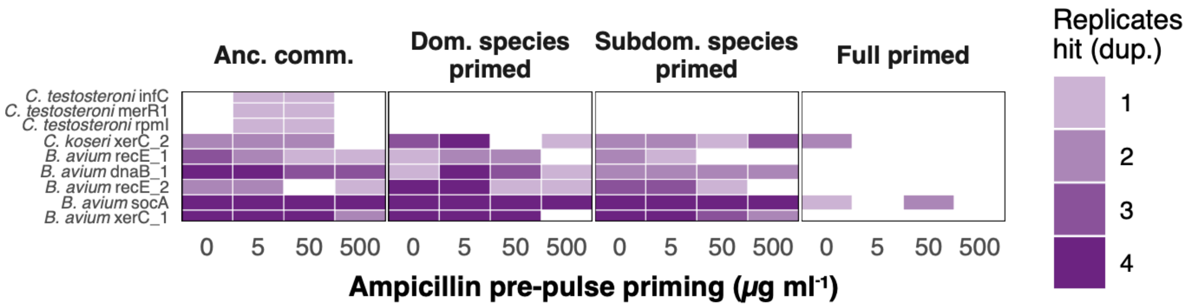


**Figure S6 | Recurrent gene duplications across community-level pre-pulse priming and species-level resistance priming backgrounds.** Heatmap of recurrent (detected in more than one independent community) **gene duplications** observed during the main antibiotic pulse, shown across ampicillin pre-pulse levels (0, 5, 50, 500 μg $mL^{-1}$; *x*-axis) and resistance priming backgrounds (facets, left to right: ancestral community; dominant species *Aeromonas caviae* resistance primed; subdominant species *Pseudomonas chlororaphis* primed; full primed community). Each tile indicates the number of independent replicate populations (1–4) in which a duplication affecting that gene was detected; darker tiles denote more replicates. Rows are grouped by species (color-coded; species legend shown below).

### Supplementary Tables

**Supplementary Table S1 | ANOVA results for linear regression model on initial strain log10 MIC values** (*N* = 23/24 strains × ancestral/pre-exposed versions = 46 strains; one strain, *Chitinophaga sancti* HAMBI 1988, lost to failure to grow in culture conditions).

| Factor | Df | Sum Sq | Mean Sq | *F*-value | *p*-value |
|---|---|---|---|---|---|
| Strain | 22 | 250.59840 | 11.39084 | 48.63886 | 8.67E-25 |
| Ampicillin pre-exposure | 1 | 6.65393 | 6.65393 | 28.41228 | 2.88E-06 |
| Strain × ampicillin pre-exposure | 22 | 34.66007 | 1.57546 | 6.72720 | 2.98E-08 |
| Residuals | 46 | 10.77284 | 0.23419 | | |

**Supplementary Table S2 | ANOVA results for linear regression model on initial strain carrying capacity (*k*) values** (*N* = 23/24 strains × ancestral/pre-exposed versions = 46 strains; one strain, *Chitinophaga sancti* HAMBI 1988, lost to failure to grow in culture conditions).

| Factor | Df | Sum Sq | Mean Sq | *F* value | *p*-value |
|---|---|---|---|---|---|
| Strain | 22 | 3.881419 | 0.176428 | 18.05484 | 7.64E-16 |
| Ampicillin pre-exposure | 1 | 0.048563 | 0.048563 | 4.969695 | 0.03072 |
| Strain x ampicillin pre-exposure | 22 | 1.137246 | 0.051693 | 5.290024 | 9.93E-07 |
| Residuals | 46 | 0.449502 | 0.009772 | | |

**Supplementary Table S3 | ANOVA results for linear regression model of community Bray–Curtis distance to the post-disturbance state (day 16 centroid), measured at day 8 (after pre-pulse).** *N* = 4 pre-exposure backgrounds × 4 pre-pulse levels × 4 replicates = 64 observations.

| Factor | Df | Sum Sq | Mean Sq | *F*-value | *p*-value |
|---|---|---|---|---|---|
| Pre-pulse ampicillin level | 1 | 0.194251 | 0.194251 | 174.41 | <0.001 |
| Pre-exposure background | 3 | 0.052042 | 0.017347 | 15.58 | <0.001 |
| Pre-pulse × background | 3 | 0.014553 | 0.004851 | 4.36 | 0.008 |
| Residuals | 56 | 0.062372 | 0.001114 | | |

**Supplementary Table S4 | ANOVA results for linear regression models of temporal roughness of Bray-Curtis distance trajectories.** $N = 4$ pre-exposure backgrounds × 4 pre-pulse levels × 4 replicates = 64 observations.

**Pre-disturbance**

| Factor | Df | Sum Sq | Mean Sq | *F*-value | *p-value* |
|---|---|---|---|---|---|
| Pre-exposure background | 3 | 0.32764 | 0.109213 | 26.40 | <0.001 |
| Residuals | 60 | 0.24822 | 0.004137 | | |

**Post-disturbance**

| Factor | Df | Sum Sq | Mean Sq | *F*-value | *p*-value |
|---|---|---|---|---|---|
| Pre-exposure background | 3 | 3.5444 | 1.18145 | 134.74 | <0.001 |
| Residuals | 60 | 0.5261 | 0.00877 | | |

**Post-recovery**

| Factor | Df | Sum Sq | Mean Sq | *F*-value | *p*-value |
|---|---|---|---|---|---|
| Pre-exposure background | 3 | 4.3433 | 1.44777 | 74.53 | <0.001 |
| Residuals | 60 | 1.1655 | 0.01942 | | |

**Post-hoc contrasts (all pre-exposed vs. each other background)**
All contrasts $p < 0.001$ in all phases.

**Supplementary Table S5 | ANOVA results for generalized least squares (GLS) regression model on community MIC (scalar product of log MIC and species relative abundance across community) in ampicillin pulse serial passage experiment.** An autocorrelation structure of order 1 was used for repeated measurements of the same community over time. ($N =$ 64 communities × 5 time points = 320 samples).

| Factor | DF | *F*-value | *p*-value |
|---|---|---|---|
| (Intercept) | 1 | 65353.0 | 0 |
| Strain pre-exposure | 3 | 933.0 | 2.23E-145 |
| Day | 4 | 270.0 | 1.02E-94 |
| Pre-pulse | 1 | 103.0 | 7.87E-21 |
| Strain pre-exposure × day | 12 | 21.4 | 3.87E-33 |
| Strain pre-exposure × pre-pulse | 3 | 0.904 | 0.440 |
| Day × pre-pulse | 4 | 59.5 | 2.71E-36 |
| Strain pre-exposure × day × pre-pulse | 12 | 3.81 | 2.15E-05 |

**Supplementary Table S6 | ANOVA results for generalized least squares (GLS) regression model on community carrying capacity *k* (scalar product of *k* and species relative abundance across community) in ampicillin pulse serial passage experiment.** An autocorrelation structure of order 1 was used for repeated measurements of the same community over time. ($N$ = 64 communities × 5 time points = 320 samples).

| Factor | DF | *F*-value | *p*-value |
|---|---|---|---|
| (Intercept) | 1 | 735446.2 | 0 |
| Strain pre-exposure | 3 | 368.782 | 6.58E-97 |
| Day | 4 | 747.5877 | 4.7E-148 |
| Pre-pulse | 1 | 12.17608 | 0.000562 |
| Strain pre-exposure x day | 12 | 49.677 | 3.25E-62 |
| Strain pre-exposure x pre-pulse | 3 | 1.177611 | 0.318566 |
| Day x pre-pulse | 4 | 17.06855 | 1.54E-12 |
| Strain pre-exposure x day x pre-pulse | 12 | 4.493744 | 1.3E-06 |

**Supplementary Table S7 | ANOVA results for linear regression model of community Bray–Curtis distance to the post-recovery state (day 20 centroid), measured at day 12 (after intermediate recovery).** $N$ = 4 pre-exposure backgrounds × 4 pre-pulse levels × 4 replicates = 64 observations.

| Factor | Df | Sum Sq | Mean Sq | *F*-value | *p*-value |
|---|---|---|---|---|---|
| Pre-pulse ampicillin level (scaled) | 1 | 0.064026 | 0.064026 | 51.61 | <0.001 |
| Pre-exposure background | 3 | 0.031828 | 0.010609 | 8.55 | <0.001 |
| Pre-pulse × background | 3 | 0.004736 | 0.001579 | 1.27 | 0.293 |
| Residuals | 56 | 0.069469 | 0.001241 | | |

**Supplementary Table S8 | ANOVA results for generalized least squares (GLS) regression model on species richness in ampicillin pulse serial passage experiment.** An autocorrelation structure of order 1 was used for repeated measurements of the same community over time. ($N$ = 64 communities × 5 time points = 320 samples).

| Factor | DF | *F*-value | *p*-value |
|---|---|---|---|
| (Intercept) | 1 | 34630.51 | 0 |
| Strain pre-exposure | 3 | 6.142028 | 0.000456 |
| Pre-pulse | 1 | 40.19106 | 8.21E-10 |
| Day | 4 | 167.23 | 8.45E-76 |
| Pre-pulse x day | 4 | 3.568737 | 0.007301 |

**Supplementary Table S9 | ANOVA results for generalized least squares (GLS) regression model on Shannon diversity in ampicillin pulse serial passage experiment.** An autocorrelation structure of order 1 was used for repeated measurements of the same community over time. (*N* = 64 communities × 5 time points = 320 samples).

| Factor | DF | *F*-value | *p*-value |
|---:|---:|---:|---:|
| (Intercept) | 1 | 89236.95 | 0 |
| Strain pre-exposure | 3 | 26.68027 | 2.69E-15 |
| Pre-pulse | 1 | 122.1578 | 5.42E-24 |
| Day | 4 | 95.87327 | 3.73E-52 |
| Strain pre-exposure x day | 12 | 19.92813 | 1.13E-31 |
| Pre-pulse x day | 4 | 44.14135 | 5.03E-29 |

**Supplementary Table S10 | ANOVA results for generalized least squares (GLS) regression model on ampicillin concentration in ampicillin pulse serial passage experiment.** An autocorrelation structure of order 1 was used for repeated measurements of the same community over time. (*N* = 64 communities × 5 time points = 320 samples).

| Factor | DF | *F*-value | *p*-value |
|---:|---:|---:|---:|
| (Intercept) | 1 | 457.5151 | 5.59E-57 |
| Strain pre-exposure | 3 | 0.500588 | 0.682234 |
| Pre-pulse | 1 | 1.476281 | 0.225582 |
| Day | 1 | 0.545411 | 0.460939 |
| Strain pre-exposure x pre-pulse | 3 | 1.464031 | 0.225062 |
| Strain pre-exposure x day | 3 | 0.529956 | 0.662144 |
| Pre-pulse x day | 1 | 17.51927 | 4.03E-05 |
| Strain pre-exposure x pre-pulse x day | 3 | 0.327285 | 0.805634 |

**Supplementary Table S11 | ANOVA results for linear regression model of change in non-*Aeromonas* fraction during the main pulse (day 16 − day 12) in the 20-day ampicillin-pulse experiment.** *N* = 4 pre-exposure backgrounds × 4 pre-pulse levels × 4 replicates = 64 observations.

| Factor | Df | Sum Sq | Mean Sq | *F*-value | *p*-value |
|---:|---:|---:|---:|---:|---:|
| Pre-exposure background | 3 | 0.4680 | 0.1560 | 179.2 | <0.001 |
| Pre-pulse ampicillin level | 3 | 0.0200 | 0.00668 | 7.67 | <0.001 |
| Pre-pulse × background | 9 | 0.0122 | 0.00135 | 1.55 | 0.157 |
| Residuals | 48 | 0.0418 | 0.000871 | | |

**Supplementary Table S12 | PERMANOVA results for mutation profiles in ampicillin pulse serial passage experiment after main pulse (day 16).** ($N$ = 64 communities).

| Factor | Df | SumOfSqs | $R^2$ | $F$ | Pr(>$F$) |
|---|---|---|---|---|---|
| Pre-pulse | 1 | 2.64908 | 0.028873 | 2.130726 | 0.0231 |
| Strain pre-exposure | 3 | 13.625 | 0.148501 | 3.652984 | 0.0001 |
| Pre-pulse x strain pre-exposure | 3 | 5.852482 | 0.063787 | 1.569103 | 0.0303 |
| Residual | 56 | 69.62344 | 0.758839 | | |
| Total | 63 | 91.75 | 1 | | |

**PERMANOVA post-hoc results for strain pre-exposure treatment.**

| Factor | Df | SumOfSqs | $R^2$ | $F$ | Pr(>$F$) |
|---|---|---|---|---|---|
| **All ancestral vs. all pre-exposed** | 1 | 8.4375 | 0.189873 | 7.03125 | 0.001 |
| Residual | 30 | 36 | 0.810127 | | |
| Total | 31 | 44.4375 | 1 | | |
| **All ancestral vs. pre-exposed 1972** | 1 | 1.625 | 0.034759 | 1.080332 | 0.388 |
| Residual | 30 | 45.125 | 0.965241 | | |
| Total | 31 | 46.75 | 1 | | |
| **All ancestral vs. pre-exposed 1977** | 1 | 0.8125 | 0.020668 | 0.633117 | 0.859 |
| Residual | 30 | 38.5 | 0.979332 | | |
| Total | 31 | 39.3125 | 1 | | |
| **All pre-exposed vs. pre-exposed 1972** | 1 | 8.125 | 0.170157 | 6.15142 | 0.001 |
| Residual | 30 | 39.625 | 0.829843 | | |
| Total | 31 | 47.75 | 1 | | |
| **All pre-exposed vs. pre-exposed 1977** | 1 | 7.25 | 0.180124 | 6.590909 | 0.001 |
| Residual | 30 | 33 | 0.819876 | | |
| Total | 31 | 40.25 | 1 | | |
| **Pre-exposed 1972 vs. pre-exposed 1977** | 1 | 1 | 0.023188 | 0.712166 | 0.812 |
| Residual | 30 | 42.125 | 0.976812 | | |
| Total | 31 | 43.125 | 1 | | |

**Test for multivariate homogeneity of group dispersions (variances) in mutation profiles in different pre-exposure histories in ampicillin pulse serial passage experiment after main pulse (day 16).** ($N$ = 64 communities).

| Df | Sum Sq | Mean Sq | $F$ value | Pr(>$F$) |
|---|---|---|---|---|
| 3 | 0.62234 | 0.207447 | 1.09257 | 0.359276 |
| 60 | 11.39223 | 0.189871 | | |

**Supplementary Table S13 | ANOVA results for Poisson regression model on mutation count in ampicillin pulse serial passage experiment after main pulse (day 16).** (*N* = 64 communities).

| Factor | Df | Deviance | Resid. Df | Resid. Dev | Pr(>Chi) |
|---|---|---|---|---|---|
| NULL | | | 63 | 49.48956 | |
| Pre-pulse | 1 | 3.724348 | 62 | 45.76521 | 0.053625 |
| Strain pre-exposure | 3 | 2.184908 | 59 | 43.58031 | 0.534927 |
| Pre-pulse x strain pre-exposure | 3 | 3.083416 | 56 | 40.49689 | 0.378942 |

### Supplementary references


1. Hogle SL, Hepolehto I, Ruokolainen L, Cairns J, Hiltunen T. Effects of phenotypic variation on consumer coexistence and prey community structure. *Ecol Lett* 2022;**25**:307–319. https://doi.org/10.1111/ele.13924
2. Cairns J, Hogle S, Alitupa E, Mustonen V, Hiltunen T. Pre-exposure of abundant species to disturbance improves resilience in microbial metacommunities. *Nat Ecol Evol* 2025;**9**:395–405. https://doi.org/10.1038/s41559-024-02624-0
3. Chen S, Zhou Y, Chen Y, Gu J. fastp: an ultra-fast all-in-one FASTQ preprocessor. *Bioinformatics* 2018;**34**:i884–i890. https://doi.org/10.1093/bioinformatics/bty560
4. Hogle SL, Tamminen M, Hiltunen T. Complete genome sequences of 30 bacterial species from a synthetic community. *Microbiol Resour Announc* 2024;**13**:e00111-24. https://doi.org/10.1128/mra.00111-24
5. Li H. Aligning sequence reads, clone sequences and assembly contigs with BWA-MEM. *arXiv* 2013; arXiv:1303.3997. https://arxiv.org/abs/1303.3997
6. Danecek P, Bonfield JK, Liddle J, Marshall J, Ohan V, Pollard MO *et al*. Twelve years of SAMtools and BCFtools. *GigaScience* 2021;**10**:giab008. https://doi.org/10.1093/gigascience/giab008
7. Benjamin D, Sato T, Cibulskis K, Getz G, Stewart C, Lichtenstein L. Calling somatic SNVs and indels with Mutect2. *bioRxiv* 2019;861054. https://doi.org/10.1101/861054
8. Cingolani P, Platts A, Wang LL, Coon M, Nguyen T, Wang L, Land SJ, Lu X, Ruden DM. A program for annotating and predicting the effects of single nucleotide polymorphisms, SnpEff: SNPs in the genome of *Drosophila melanogaster* strain w1118; iso-2; iso-3. *Fly (Austin)* 2012;**6**:80–92. https://doi.org/10.4161/fly.19695
9. Cingolani P, Patel VM, Coon M, Nguyen T, Land SJ, Ruden DM *et al*. Using *Drosophila melanogaster* as a model for genotoxic chemical mutational studies with a new program, SnpSift. *Front. Genet.* 2012;**3**:35. https://doi.org/10.3389/fgene.2012.00035
10. Bushnell B. BBMap: A fast, accurate, splice-aware aligner. Lawrence Berkeley National Laboratory. *LBNL Report* 2014;LBNL-7065E. https://escholarship.org/uc/item/1h3515gn
11. Jugas R, Sedlar K, Vitek M, Nykrynova M, Barton V, Bezdicek M *et al*. CNproScan: hybrid CNV detection for bacterial genomes. *Genomics* 2021;**113**:3103–3111. https://doi.org/10.1016/j.ygeno.2021.06.040
12. Pockrandt C, Alzamel M, Iliopoulos CS, Reinert K. GenMap: ultra-fast computation of genome mappability. *Bioinformatics* 2020;**36**:3687–3692. https://doi.org/10.1093/bioinformatics/btaa222
13. Patel H, Ewels P, Peltzer A, Manning J, Botvinnik O, Sturm G *et al*. nf-core/rnaseq: nf-core/rnaseq v3.14.0 – Hassium Honey Badger. Zenodo 2024. https://doi.org/10.5281/zenodo.10471647
14. Ewels PA, Peltzer A, Fillinger S, Patel H, Alneberg J, Wilm A *et al*. The nf-core framework for community-curated bioinformatics pipelines. *Nat Biotechnol* 2020;**38**:276–278. https://doi.org/10.1038/s41587-020-0439-x
15. Patro R, Duggal G, Love MI, Irizarry RA, Kingsford C. Salmon provides fast and bias-aware quantification of transcript expression. *Nat Methods* 2017;**14**:417–419. https://doi.org/10.1038/nmeth.4197

16. Hogle SL, Ruusulehto L, Cairns J, Hultman J, Hiltunen T. Localized coevolution between microbial predator and prey alters community-wide gene expression and ecosystem function. *ISME J* 2023;**17**:514–524. https://doi.org/10.1038/s41396-023-01361-9
17. Pantano L. DEGreport: Report of DEG Analysis. R package version 1.40.1, Bioconductor 2024. http://lpantano.github.io/DEGreport/
18. Daily K, Ho Sui SJ, Schriml LM, Dexheimer PJ, Salomonis N, Schroll R *et al*. Molecular, phenotypic, and sample-associated data to describe pluripotent stem cell lines and derivatives. *Sci. Data* 2017;**4**:170030. https://doi.org/10.1038/sdata.2017.30
19. Love MI, Huber W, Anders S. Moderated estimation of fold change and dispersion for RNA-seq data with DESeq2. *Genome Biol* 2014;**15**:550. https://doi.org/10.1186/s13059-014-0550-8
20. Huerta-Cepas J, Forslund K, Coelho LP, Szklarczyk D, Jensen LJ, von Mering C *et al*. Fast genome-wide functional annotation through orthology assignment by eggNOG-mapper. *Mol. Biol. Evol.* 2017;**34**:2115–2122. https://doi.org/10.1093/molbev/msx148
21. Huerta-Cepas J, Szklarczyk D, Heller D, Hernández-Plaza A, Forslund SK, Cook H *et al.* eggNOG 5.0: a hierarchical, functionally and phylogenetically annotated orthology resource based on 5090 organisms and 2502 viruses. *Nucleic Acids Res.* 2019;**47**:D309–D314. https://doi.org/10.1093/nar/gky1085
22. Alcock BP, Huynh W, Chalil R, Smith KW, Raphenya AR, Wlodarski MA *et al*. CARD 2023: expanded curation, support for machine learning, and resistome prediction at the Comprehensive Antibiotic Resistance Database. *Nucleic Acids Res.* 2023;**51**:D690–D699. https://doi.org/10.1093/nar/gkac920
23. Wu T, Hu E, Xu S, Chen M, Guo P, Dai Z *et al*. clusterProfiler 4.0: a universal enrichment tool for interpreting omics data. *Innovation (Camb.)* 2021;**2**:100141. https://doi.org/10.1016/j.xinn.2021.100141
24. Glen KA, Lamont IL. Penicillin-binding protein 3 sequence variations reduce susceptibility of *Pseudomonas aeruginosa* to β-lactams but inhibit cell division. *J Antimicrob Chemother* 2024;**79**:2170–2178. https://doi.org/10.1093/jac/dkae203
25. Hugonnet JE, Mengin-Lecreulx D, Monton A, den Blaauwen T, Carbonnelle E, Veckerlé C *et al*. Factors essential for L,D-transpeptidase-mediated peptidoglycan cross-linking and β-lactam resistance in *Escherichia coli*. *eLife* 2016;**5**:e19469. https://doi.org/10.7554/eLife.19469
26. Gross R, Yelin I, Lázár V, Datta MS, Kishony R. Beta-lactamase dependent and independent evolutionary paths to high-level ampicillin resistance. *Nat Commun* 2024;**15**:5383. https://doi.org/10.1038/s41467-024-49621-2
27. Mittal S, Mallik S, Sharma S, Virdi JS. Characteristics of β-lactamases and their genes (*blaA* and *blaB*) in *Yersinia intermedia* and *Y. frederiksenii*. *BMC Microbiol* 2007;**7**:25. https://doi.org/10.1186/1471-2180-7-25
28. Garoff L, Huseby DL, Praski Alzrigat L, Hughes D. Effect of aminoacyl-tRNA synthetase mutations on susceptibility to ciprofloxacin in *Escherichia coli*. *J Antimicrob Chemother* 2018;**73**:3285–3292. https://doi.org/10.1093/jac/dky356
29. Yang HD, Jeong H, Kim Y, Lee HS. The *cysS* gene (ncgl0127) of *Corynebacterium glutamicum* is required for sulfur assimilation and affects oxidative stress-responsive cysteine import. *Res Microbiol* 2022;**173**:103983. https://doi.org/10.1016/j.resmic.2022.103983

30. Dordel J, Kim C, Chung M, de la Gándara MP, Holden MTJ, Parkhill J *et al*. Novel determinants of antibiotic resistance: identification of mutated loci in highly methicillin-resistant subpopulations of methicillin-resistant *Staphylococcus aureus*. *mBio* 2014;**5**:e01000-13. https://doi.org/10.1128/mbio.01000-13
31. Cam K, Rome G, Krisch HM, Bouché JP. RNase E processing of essential cell division genes mRNA in *Escherichia coli*. *Nucleic Acids Res* 1996;**24**:3065–3070. https://doi.org/10.1093/nar/24.15.3065
32. Ochoa-Sánchez LE, Martínez JL, Gil-Gil T. Evolution of resistance against ciprofloxacin, tobramycin, and trimethoprim/sulfamethoxazole in the environmental opportunistic pathogen *Stenotrophomonas maltophilia*. *Antibiotics* 2024;**13**:330. https://doi.org/10.3390/antibiotics13040330
33. Balasubramanian D, Kumari H, Mathee K. *Pseudomonas aeruginosa* AmpR: an acute–chronic switch regulator. *Pathog Dis* 2015;**73**:1–14. https://doi.org/10.1111/2049-632X.12208
34. Azimi A, Aslanimehr M, Yaseri M, Shadkam M, Douraghi M. Distribution of *smf-1*, *rmlA*, *spgM* and *rpfF* genes among *Stenotrophomonas maltophilia* isolates in relation to biofilm-forming capacity. *J Glob Antimicrob Resist* 2020;**23**:321–326. https://doi.org/10.1016/j.jgar.2020.10.011
35. Hoffmann J, Hogle S, Hiltunen T, Becks L. Temporal changes in the role of species sorting and evolution determine community dynamics. *Ecol Lett* 2024;**28**:e70033. https://doi.org/10.1111/ele.70033
36. Bos J, Zhang Q, Vyawahare S, Rogers E, Rosenberg SM, Austin RH. Emergence of antibiotic resistance from multinucleated bacterial filaments. *Proc Natl Acad Sci USA* 2015;**112**:178–183. https://doi.org/10.1073/pnas.1420702111
37. Hong J, Li X, Jiang M, Hong R. Co-expression mechanism analysis of different tachyplesin I–resistant strains in *Pseudomonas aeruginosa* based on transcriptome sequencing. *Front Microbiol* 2022;**13**:871290. https://doi.org/10.3389/fmicb.2022.871290
38. Andersson DI, Nicoloff H, Hjort K. Mechanisms and clinical relevance of bacterial heteroresistance. *Nat Rev Microbiol* 2019;**17**:479–496. https://doi.org/10.1038/s41579-019-0218-1
39. Cianciulli Sesso A, Lilić B, Amman F, Wolfinger MT, Sonnleitner E, Bläsi U. Gene expression profiling of *Pseudomonas aeruginosa* upon exposure to colistin and tobramycin. *Front Microbiol* 2021;**12**:626715. https://doi.org/10.3389/fmicb.2021.626715
40. Ledger EVK, Lau K, Tate EW, Edwards AM. XerC is required for the repair of antibiotic- and immune-mediated DNA damage in *Staphylococcus aureus*. *Antimicrob. Agents Chemother.* 2023;**67**:e01206-22. https://doi.org/10.1128/aac.01206-22
